# Mitochondrial-Derived Compartments Facilitate Cellular Adaptation to Amino Acid Stress

**DOI:** 10.1101/2020.03.13.991091

**Authors:** Max-Hinderk Schuler, Alyssa M. English, Thane J. Campbell, Janet M. Shaw, Adam L. Hughes

**Author notes:** These authors contributed equally to this work (Max-Hinderk Schuler, Alyssa M. English). Correspondence: Department of Biochemistry, University of Utah School of Medicine, 15 N. Medical Drive East, RM 4100, Salt Lake City, UT, 84112.

## Abstract

Amino acids are essential building blocks of life. However, increasing evidence suggests that elevated amino acids cause cellular toxicity associated with numerous metabolic disorders. How cells cope with elevated amino acids remains poorly understood. Here, we show that a previously identified cellular structure, the <u>m</u>itochondrial-<u>d</u>erived <u>c</u>ompartment (MDC), is a dynamic, lumen-containing organelle that functions to protect cells from amino acid stress. In response to amino acid elevation, MDCs are generated from mitochondria, where they selectively sequester and remove Tom70, a surface receptor required for import of nutrient carriers of the SLC25 family. MDC formation is regulated by levels of mitochondrial carriers, and its activation by amino acids occurs simultaneously with removal of plasma membrane-localized transporters via the multi-vesicular body (MVB) pathway. Combined loss of MDC and MVB formation renders cells sensitive to elevated amino acids, suggesting these pathways operate as a coordinated network to protect cells from amino acid toxicity.

## INTRODUCTION

Amino acids are essential metabolites utilized as fuel sources, signaling molecules, and precursors for the biosynthesis of proteins, lipids, heme, nucleotides, and other cellular molecules (Ljungdahl and Daignan-Fornier, 2012). As with most metabolites, cells must maintain amino acid homeostasis at all times, and they are equipped with numerous systems that monitor amino acid concentrations and adjust the rates of amino acid acquisition, storage, and utilization accordingly (Efeyan et al., 2015). To date, we know a great deal about the impact of amino acid starvation on cells, as well as the signaling pathways and remodeling systems such as autophagy that operate to maintain cellular health when amino acids are in short supply (Efeyan et al., 2015; Rabinowitz and White, 2010). On the other hand, elevated amino acids can lead to cellular toxicity and are associated with aging and numerous metabolic disorders, including insulin resistance and a host of inborn errors of amino acid metabolism (Aliu et al., 2018; Newgard et al., 2009; Ruiz et al., 2017; Soultoukis and Partridge, 2016). In contrast to our knowledge of cellular adaptation to amino acid starvation, we understand little about the mechanisms that drive amino acid toxicity and pathways that protect cells from amino acid overabundance stress (Wellen and Thompson, 2010).

We recently discovered that the yeast lysosome (vacuole) functions as a safeguard against cellular amino acid toxicity through its ability to import and sequester amino acids (Hughes and Gottschling, 2012; Hughes et al., 2020). Defects in vacuolar amino acid compartmentation impair mitochondrial respiration and negatively impact cellular health (Hughes and Gottschling, 2012; Hughes et al., 2020). In addition, prior work in yeast has shown that the regulation of plasma membrane (PM)-localized nutrient transporters via the multi-vesicular body (MVB) pathway also serves as a key mechanism to control cellular nutrient uptake, and protects cells from amino acid toxicity (Katzmann et al., 2002; Risinger et al., 2006; Rubio-Texeira and Kaiser, 2006; Ruiz et al., 2017). Beyond lysosomes and MVBs, additional mechanisms cells utilize to protect themselves from amino acid toxicity remain unclear.

While investigating the impact of lysosome failure on mitochondrial health, we identified a new cellular structure that forms from mitochondria when lysosomal acidification is impaired, called the <u>m</u>itochondrial-<u>d</u>erived <u>c</u>ompartment (MDC) (Hughes et al., 2016). Upon formation, MDCs selectively incorporate a number of mitochondrial proteins including Tom70, an outer membrane (OM) import receptor for mitochondrial nutrient transporters (Sollner et al., 1990). By contrast, MDCs exclude most other mitochondrial proteins, including those in the mitochondrial matrix, the intermembrane space (IMS), and the majority of inner membrane (IM) proteins. After formation, MDCs are released from mitochondria via mitochondrial fission and are degraded by autophagy (Hughes et al., 2016). Currently, we know that MDCs are Tom70-enriched foci that associate with mitochondria when cells lose lysosome acidification. Beyond that, we understand little about the dynamics and regulation of MDC formation, as well as the function of this new cellular compartment.

Here, we show that MDCs are dynamic mitochondrial-associated compartments that are generated in response to perturbations in intracellular amino acid homeostasis. Specifically, we find that high levels of branched-chain amino acids (BCAAs) and their breakdown products promote MDC formation. Our data indicate that MDCs sequester Tom70 away from mitochondria and might thereby provide cells with a mechanism to fine-tune levels of mitochondrial nutrient transporters in response to changes in cellular amino acid load. Consistent with this idea, increasing nutrient transporter levels on mitochondria via Tom70 or nutrient transporter overexpression triggers constitutive MDC formation. Finally, we show that MDC formation and removal of nutrient transporters from the PM via the MVB pathway are activated in parallel, and that combined loss of both systems impairs the cell’s ability to cope with toxic levels of BCAAs. Overall, our data suggest that MDCs are part of a coordinated cellular program that operates to protect cells from amino acid stress.

## RESULTS

### MDCs are dynamic, lumen-containing organelles that stably associate with mitochondria

We previously identified MDCs in budding yeast, *S. cerevisiae*, as bright foci that form in association with mitochondria during aging or in response to acute pharmacological disruption of lysosome acidification (Hughes et al., 2016). We confirmed our prior observations by showing that treatment of cells with concanamycin A (concA), a specific inhibitor of the evolutionarily conserved Vacuolar H^+^-ATPase (V-ATPase) proton pump, triggered formation of foci containing C-terminally GFP tagged Tom70 (Figure 1A). Line-scan analysis demonstrated that MDCs exhibited an enrichment of Tom70 compared to the rest of the mitochondrial tubule and excluded the IM protein Tim50-mCherry (Yamamoto et al., 2002), a subunit of the TIM23 complex that we previously showed was excluded from MDCs (Figure 1A). Disruption of lysosome acidification via auxin-dependent depletion of the V-ATPase subunit Vma2 also induced MDC formation, indicating that MDCs form in direct response to loss of lysosome acidification (Figures S1A-D). Utilizing super-resolution microscopy, we found that MDCs are organelle-like structures with distinct lumens that reach diameters approaching ∼1 μm (Figures 1B-C). Serial optical-plane sectioning indicated that MDCs exhibited reduced diameters near the top and bottom and appeared closed at both ends, suggesting they are membrane-bound compartments (Figure 1D). In support of this latter conclusion, we previously showed that MDCs are released from mitochondria by the fission GTPase Dnm1, which acts on membrane-bound organelles (Hughes et al., 2016). MDCs excluded the mitochondrial DNA stain 4′,6-diamidino-2-phenylindole (DAPI), and did not incorporate the mitochondrial inner membrane potential-dependent dye tetramethylrhodamine (TMRM), indicating that MDCs do not contain mitochondrial DNA or a membrane potential and are thus distinct structures compared to mitochondria (Figures 1E-F).

**Figure 1.**
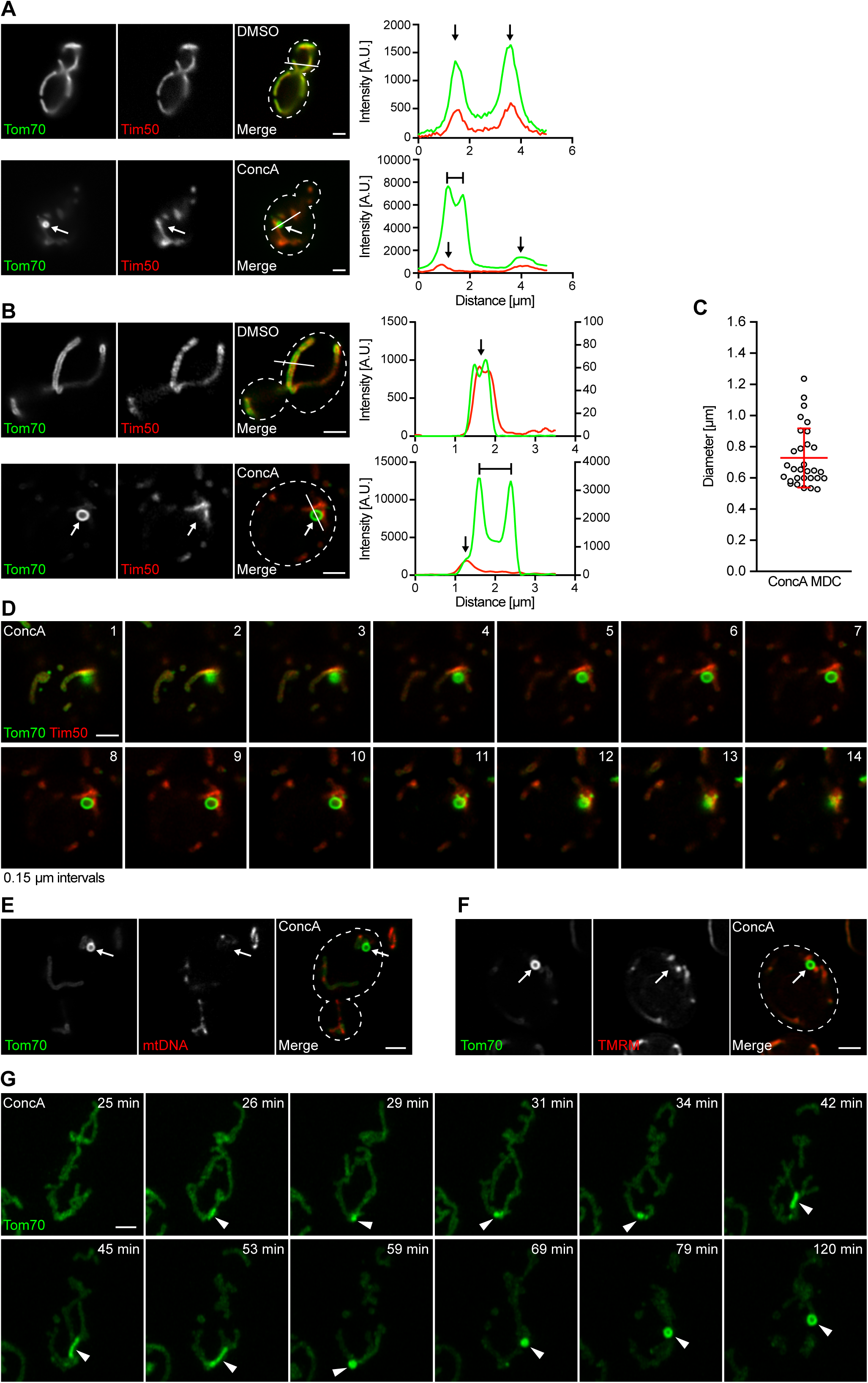
MDCs are Dynamic, Mitochondria-Associated Structures with Distinct Lumens (A) Widefield images (left) and line-scan analysis (right) of concanamycin A (ConcA)-induced MDC formation in yeast cells expressing Tom70-GFP and Tim50-mCherry. White arrow marks MDC. White line marks fluorescence intensity profile position. Line-scan Y axis corresponds to Tom70-GFP and Tim50-mCherry fluorescence intensity. Black arrow marks mitochondrial tubule. Bracket marks MDC. Scale bar = 2 µm. (B) Super-resolution images (left) and line-scan analysis (right) of ConcA-induced MDC formation in yeast cells expressing Tom70-GFP and Tim50-mCherry. White arrow marks MDC. White line marks fluorescence intensity profile position. Left and right line-scan Y axis correspond to Tom70-GFP and Tim50-mCherry fluorescence intensity, respectively. Black arrow marks mitochondrial tubule. Bracket marks MDC. Scale bar = 2 µm. (C) Scatter plot showing the diameter of ConcA-induced MDCs. Error bars show mean (0.7282 µm) ± SD of *n* = 30 MDCs. (D) Z-series corresponding to yeast cell treated with ConcA in (B). Image intervals are 0.15 µm. Scale bar = 2 µm. (E) Super-resolution images of ConcA-treated yeast cells expressing Tom70-GFP, stained with DAPI to label mitochondrial DNA. White arrow marks MDC. Scale bar = 2 µm. (F) Super-resolution images of ConcA treated yeast cells expressing Tom70-GFP stained with fluorescent membrane potential indicator TMRM. White arrow marks MDC. Scale bar = 2 µm. (G) Time-lapse images of ConcA-induced MDC formation in yeast cells expressing Tom70-GFP. Images were acquired over 120 minutes (min). Arrowhead marks MDC. Scale bar = 2 µm. See also Video S1. See also Figure S1.

To elucidate the kinetics and dynamics of MDC formation, we utilized super-resolution time-lapse imaging to visualize MDC formation over a two-hour time period. As shown in Figure 1G and Video S1, MDCs first appeared as small foci with an enrichment of Tom70-GFP compared to the rest of the tubule, usually within 20-30 minutes of concA addition (Figure 1G, 26 and 29 minute panels). MDCs remained stably associated with the mitochondrial tubule and grew in size over the 2-hour time-course, with large lumens becoming visible at late stages of formation (Figure 1G and Video S1). During and after formation, MDCs exhibited dynamic properties including frequent elongation and tubulation (Figures 1G, S1E and Video S2). Tubulation often preceded the appearance of a clear lumen, suggesting this may be a general feature of MDC formation (Figures 1G and S1E). Altogether, these data indicate that MDCs are membrane-bound, dynamic organelles capable of stable mitochondrial association.

### MDC formation is triggered by elevated cellular amino acid load

To elucidate the function of MDCs, we sought to identify the signal originating from dysfunctional lysosomes that activates MDC formation. We and others previously showed that mitochondria and lysosomes are functionally linked, and that loss of lysosome acidification impairs mitochondrial respiration (Chen et al., 2020; Dimmer et al., 2002; Hughes and Gottschling, 2012; Hughes et al., 2020; Merz and Westermann, 2009; Ohya et al., 1991; Weber et al., 2020; Yambire et al., 2019). In a recent study, we demonstrated that lysosomes support mitochondrial respiration by spatially compartmentalizing amino acids (Hughes et al., 2020). Loss of V-ATPase function prevents proper storage of amino acids in lysosomes, and causes acute amino acid stress (Figure 2A). This amino acid stress is associated with global cellular rewiring at the mRNA level, with cells turning-down amino acid biosynthesis and upregulating pathways that break down amino acids (Hughes et al., 2020). Furthermore, elevated amino acids cause mitochondrial depolarization by altering the bioavailability of intracellular iron, through an oxidant-based mechanism (Hughes et al., 2020).

**Figure 2.**
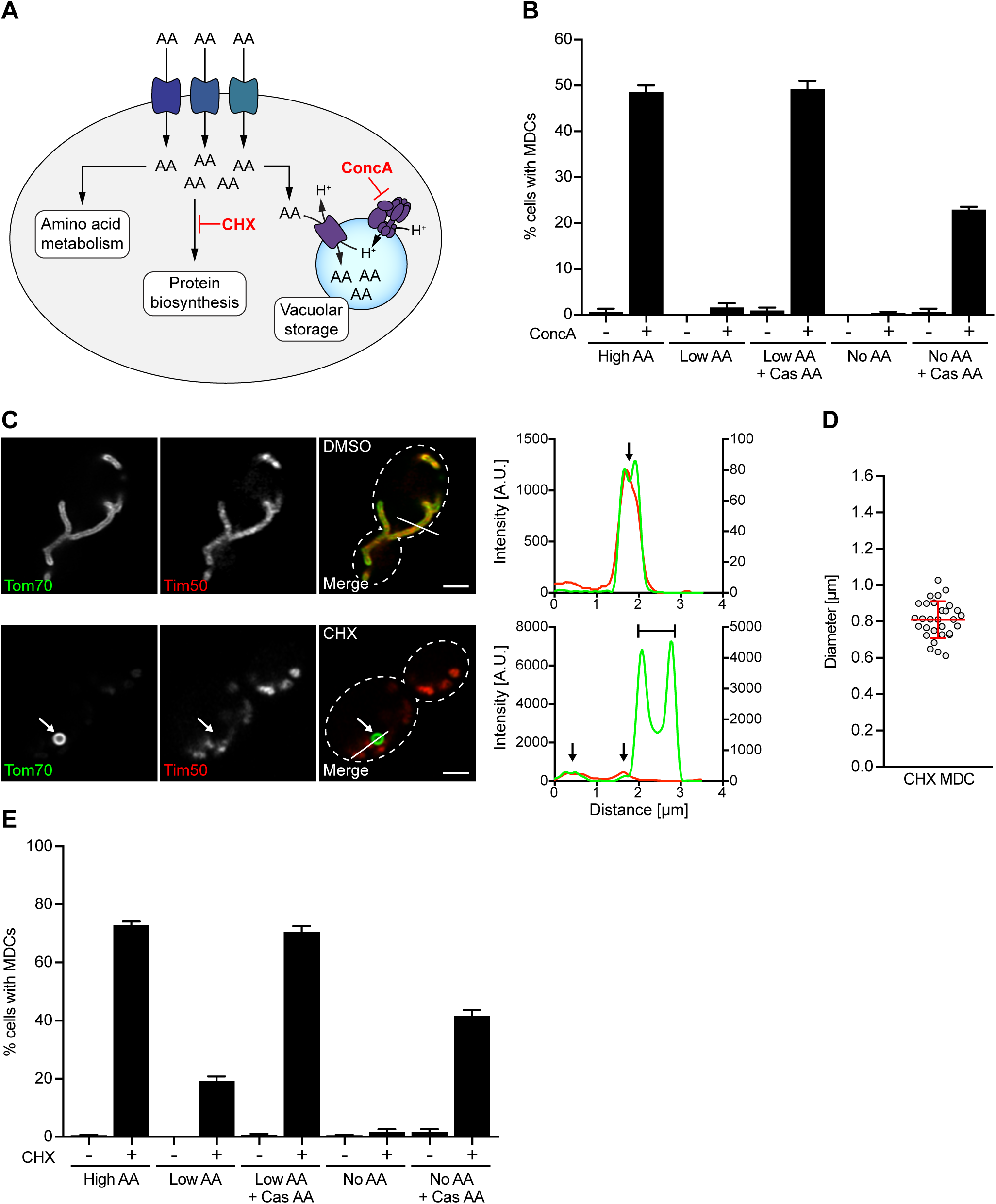
MDC Formation is Triggered in Response to Amino Acid Elevation. (A) Model of intracellular amino acid (AA) homeostasis. AAs are metabolized, utilized for protein biosynthesis or stored in the yeast vacuole (lysosome). Defects in these processes cause buildup of amino acids in the cytoplasm. (B) Quantification of concanamycin A (ConcA)-induced MDC formation in media containing high, low or no amino acids (AA). Cas AA indicates where bulk amino acids were added back to media in the form of hydrolyzed casein. Error bars show mean ± SE of three replicates with *n* = 100 cells per replicate. (C) Super-resolution images (left) and line-scan analysis (right) of cycloheximide (CHX)-induced MDC formation in yeast cells expressing Tom70-GFP and Tim50-mCherry. White arrow marks MDC. White line marks fluorescence intensity profile position. Left and right Y line-scan axis correspond to Tom70-GFP and Tim50-mCherry fluorescence intensity, respectively. Black arrow marks mitochondrial tubule. Bracket marks MDC. Scale bar = 2 µm. (D) Scatter plot showing the diameter of CHX-induced MDCs. Error bars show mean (0.8095 µm) ± SD of *n* = 30 MDCs. (E) Quantification of CHX-induced MDC formation in media containing high, low or no amino acids (AA). Cas AA indicates where bulk amino acids were added back to media in the form of hydrolyzed casein. Error bars show mean ± SE of three replicates with *n* = 100 cells per replicate. See also Figure S2.

Based on these prior observations, we reasoned that mitochondrial depolarization, iron deprivation, oxidative stress, or alterations in amino acid pools in V-ATPase-impaired cells may activate MDC formation. Treatment with the ATP synthase inhibitor oligomycin, mitochondrial depolarizing agents antimycin A and FCCP, hypoxia mimetic CoCl_2_, the ER stress inducer tunicamycin, or the oxidant hydrogen peroxide did not induce MDC formation (Figure S2A). Likewise, MDCs were not induced when cells were treated with the iron chelator BPS (Figure S2B), and iron addition to concA-treated cells, which is sufficient to restore mitochondrial respiration in the absence of a functioning vacuole (Chen et al., 2020; Hughes et al., 2020), did not inhibit concA-induced MDC formation (Figure S2B). Together, these results suggest that MDCs are not responsive to typical mitochondrial stressors, including loss of mitochondrial membrane potential, oxidative stress, ATP depletion, or iron deprivation.

Consistent with a role for amino acids in triggering MDC formation, inhibiting the V-ATPase with concA activated MDC formation when cells were cultured in rich medium containing high levels of amino acids (AA), but not in synthetic medium (Low AA), or in minimal medium completely lacking amino acids (No AA) (Figure 2B). Steady-state amino acid analysis indicated that cells grown under the latter two conditions contained lower intracellular amino acid pools when compared to cells grown in rich media (Figure S2C). Supplementation of cells with casamino acids (Cas AA) restored MDC formation in medium containing low or no amino acids (Figure 2B).

As an alternative mechanism to create acute, intracellular amino acid surplus, we treated cells with cycloheximide (CHX), which blocks incorporation of amino acids into proteins (Beugnet et al., 2003) (Figure 2A). Like concA, CHX also activated formation of Tom70-positive, Tim50-negative structures in a high percentage of cells (Figure 2C). These structures were cargo selective, dynamic, and approached sizes close to 1 μm in diameter (Figures 2C-D, S2D and Video S3). CHX-induced MDCs lacked mtDNA and membrane potential (Figures S2E-F). Importantly, depletion of amino acids prevented CHX-induced MDC formation, and supplementation with casamino acids restored MDCs, indicating that MDC induction by CHX was amino acid-dependent, and not simply due to inhibition of protein synthesis (Figure 2E). Thus, MDC formation is triggered by acute elevation of intracellular amino acids.

### BCAAs and their catabolites activate MDC formation

Next, we sought to identify the specific amino acids that trigger MDC activation. We found that addition of single amino acids to low amino acid medium restored MDC formation in the presence of concA to different extents (Figure 3A). Leucine was the most potent activator of MDC formation, followed by methionine, isoleucine, and glutamate, while several other amino acids, including arginine and proline, did not activate formation of MDCs (Figure 3A). In yeast, the most potent activators of MDC formation, including the BCAAs leucine and isoleucine as well as methionine, are all metabolized via the Ehrlich pathway, which converts amino acids to their corresponding *α*-keto acid via transamination, followed by decarboxylation to an aldehyde and oxidation to an alcohol (Figure 3B) (Hazelwood et al., 2008). To test whether downstream metabolic products of leucine catabolism also trigger MDC formation, we compared MDC formation induced by leucine to its downstream catabolic products ketoisocaproic acid (KIC), isovaleraldehye (IVA), and isoamylalcohol (IAA) (Figure 3C). Addition of KIC and IVA to low amino acid medium restored MDC formation in the presence of concA to the same extent or even more robustly than leucine, while IAA had no effect (Figure 3C). Remarkably, μM concentrations of IVA alone, without impairment of V-ATPase function or inhibition of protein synthesis, potently activated MDC formation (Figure 3D-E). Like concA and CHX, IVA-activated MDCs were cargo selective, reached sizes of nearly 1 μm in diameter, and lacked mtDNA and a membrane potential (Figures 3E-F, S3A-B). Similar results were obtained with the aldehyde derivative of methionine, which stimulated MDC formation both in the presence and absence of lysosome impairment (Figures S3C-H). Altogether, these results suggest that BCAAs and their catabolic derivatives are potent activators of MDC formation.

**Figure 3.**
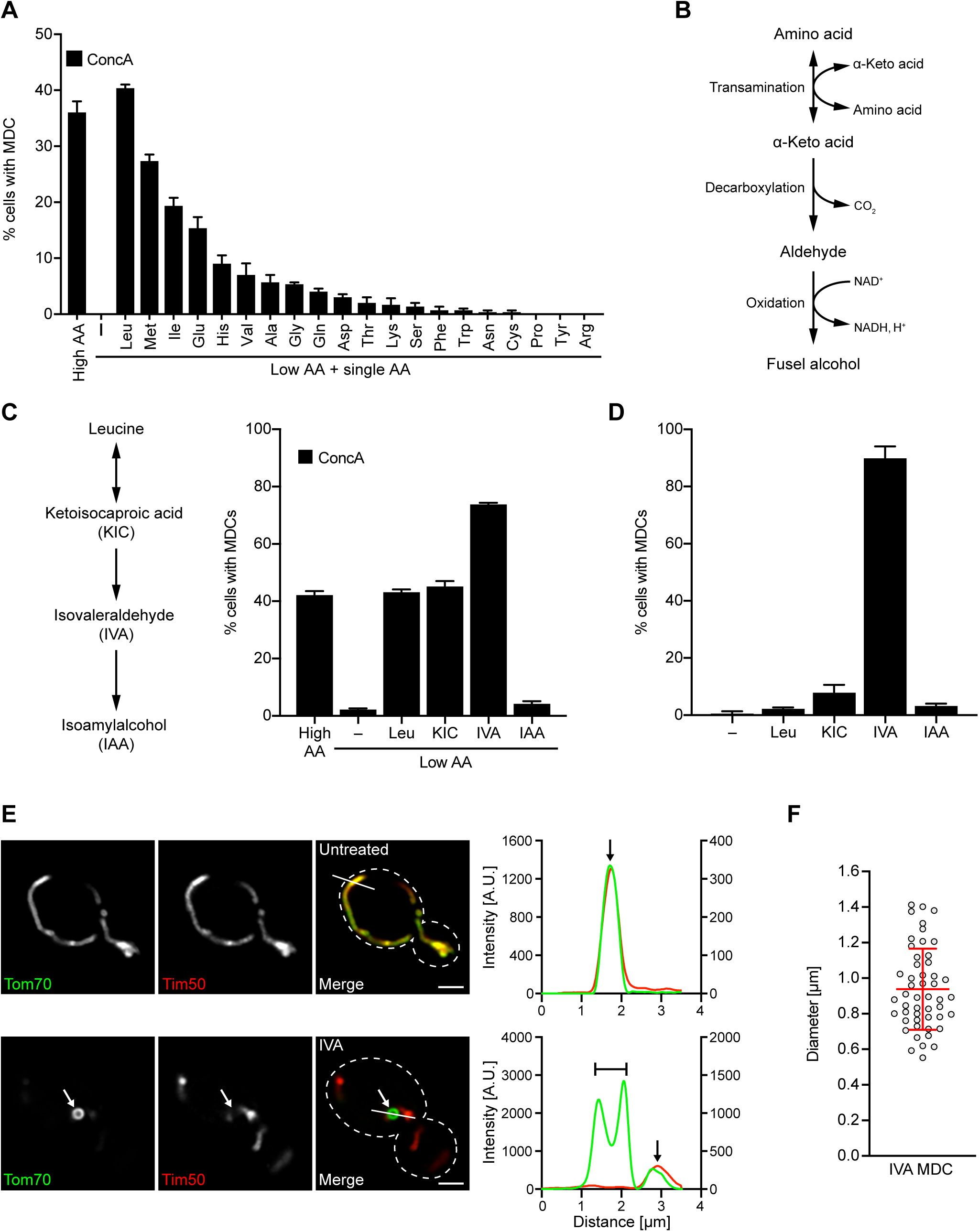
Branched-Chain Amino Acids and Their Derivatives Activate MDC Formation. (A) Quantification of concanamycin A (ConcA)-induced MDC formation in high or low amino acid medium supplemented with the indicated amino acid (AA). Error bars show mean ± SE of three replicates with *n* = 100 cells per replicate. (B) Diagram of branched chain amino acid catabolism via the Ehrlich Pathway. (C) Quantification of ConcA-induced MDC formation in low amino acid medium supplemented with leucine (Leu) or its catabolites ketoisocaproic acid (KIC), isovaleraldehyde (IVA) or isoamylalcohol (IAA). Error bars show mean ± SE of three replicates with *n* = 100 cells per replicate. (D) Quantification of MDC formation in high amino acid medium supplemented with Leu, KIC, IVA or IAA in the absence of drug treatment. Error bars show mean ± SE of three replicates with *n* = 100 cells per replicate. (E) Super-resolution images (left) and line-scan analysis (right) of IVA-induced MDC formation in yeast cells expressing Tom70-GFP and Tim50-mCherry. White arrow marks MDC. White line marks fluorescence intensity profile position. Left and right line-scan Y axis correspond to Tom70-GFP and Tim50-mCherry fluorescence intensity, respectively. Black arrow marks mitochondrial tubule. Bracket marks MDC. Scale bar = 2 µm. (F) Scatter plot showing the diameter of IVA-induced MDCs. Error bars show mean (0.938 µm) ± SD of *n* = 50 MDCs. See also Figure S3.

### Amino acids stimulate MDC formation independent of known nutrient sensing pathways

In yeast, amino acid availability is transmitted through several nutrient sensing pathways including the mechanistic Target of Rapamycin (mTOR) kinase, a central component of two independent nutrient sensing complexes, mTORC1 and mTORC2 (Saxton and Sabatini, 2017) (Figure 4A), *GCN2* (Hinnebusch, 2005), Gap1/Protein Kinase A (PKA) (Donaton et al., 2003) and the Ssy1-Ptr3-Ssy5 (SPS) sensing pathway (Ljungdahl, 2009). These pathways are activated by elevated nutrients and regulate numerous cellular processes to coordinate cell growth with nutrient availability. Since leucine is critical for mTOR signaling (Hara et al., 1998), we reasoned that increased mTOR activity in response to amino acid elevation might provide the signal to activate MDC formation (Figure 4A). To our surprise, inhibiting mTOR with rapamycin (Saxton and Sabatini, 2017) did not block concA or CHX-induced MDC formation (Figure 4B). In fact, rapamycin enhanced concA-induced MDC formation (Figure 4B), and activated cargo selective MDC formation on its own with characteristics identical to concA and CHX-induced MDC formation (Figures 4C-D, S4A-C and Video S4). Similar results were observed with Torin1 (Thoreen et al., 2009), another mTOR inhibitor (Figures S4D-E).

**Figure 4.**
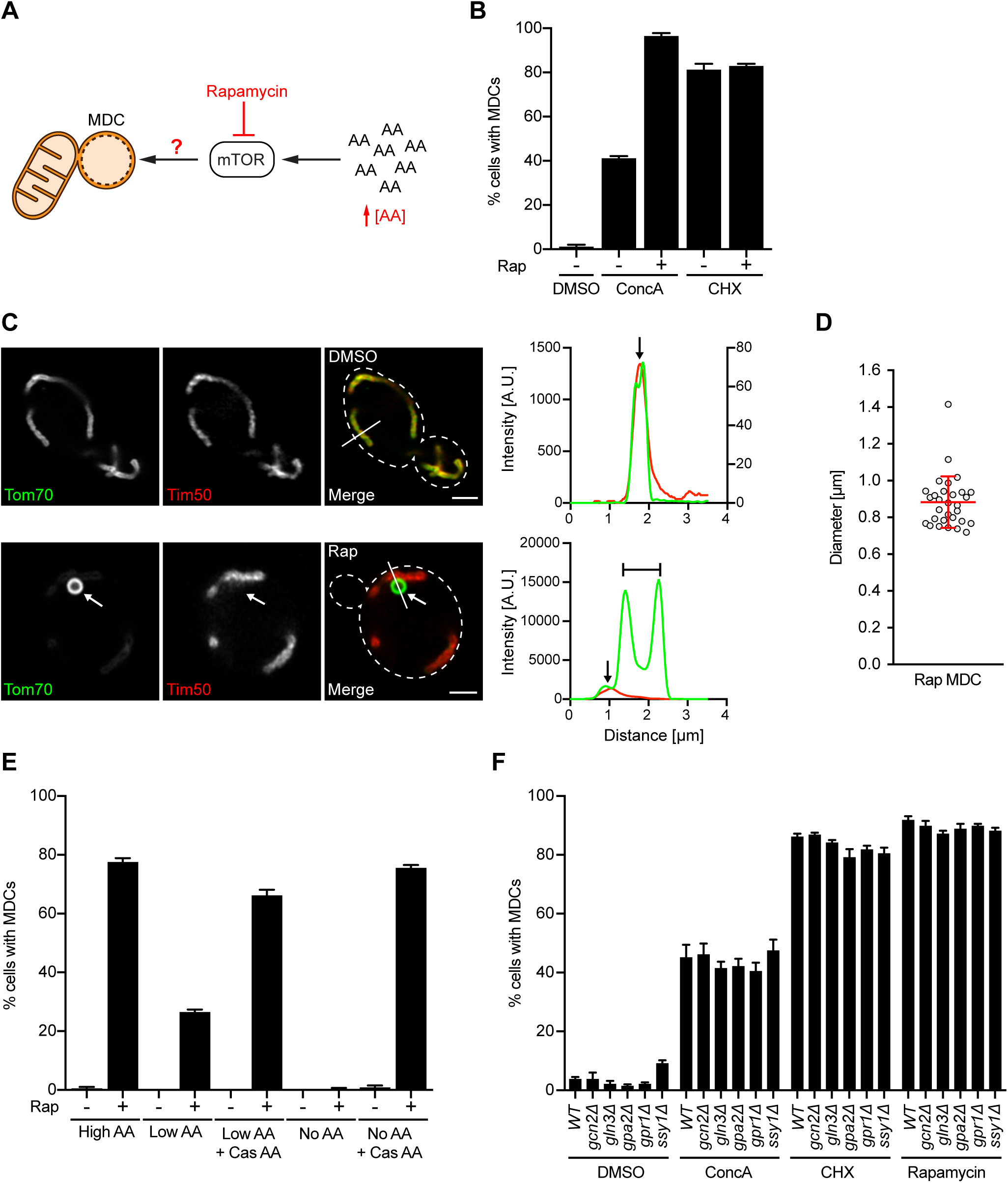
Amino Acids Activate MDC Formation Independent of Known Nutrient Sensing Pathways. (A) Schematic of amino acid (AA)-induced activation of mTOR signaling. (B) Quantification of concanamycin A (ConcA) and cycloheximide (CHX)-induced MDC formation in the absence or presence of rapamycin (Rap). Error bars show mean ± SE of three replicates with *n* = 100 cells per replicate. (C) Super-resolution images (left) and line-scan analysis (right) of Rap-induced MDC formation in yeast cells expressing Tom70-GFP and Tim50-mCherry. White arrow marks MDC. White line marks fluorescence intensity profile position. For DMSO, left and right line-scan Y axis correspond to Tom70-GFP and Tim50-mCherry fluorescence intensity, respectively. For Rap, left line-scan Y axis corresponds to both Tom70-GFP and Tim50-mCherry fluorescence intensity. Black arrow marks mitochondrial tubule. Bracket marks MDC. Scale bar = 2 µm. (D) Scatter plot showing the diameter of Rap-induced MDCs. Error bars show mean (0.8829 µm) ± SD of *n* = 30 MDCs. (E) Quantification of Rap-induced MDC formation in media containing high, low or no amino acids (AA). Cas AA indicates where bulk amino acids were added back to media in the form of hydrolyzed casein. Error bars show mean ± SE of three replicates with *n* = 100 cells per replicate. (F) Quantification of ConcA, CHX and Rap-induced MDC formation in yeast strains deficient for the indicated nutrient sensing pathway. Error bars show mean ± SE of three replicates with *n* = 100 cells per replicate. See also Figure S4.

In yeast, inhibition of mTOR signaling, which occurs naturally during amino acid starvation, enhances nutrient availability by blocking protein translation (Barbet et al., 1996), stimulating expression of the general amino acid permease Gap1 (Cardenas et al., 1999), and activating autophagy (Noda and Ohsumi, 1998). Therefore, inhibiting the mTOR pathway in nutrient-replete conditions resulted in intracellular amino acid overload, consistent with prior observations (Figure S4F) (Chen and Kaiser, 2003). As with concA and CHX, rapamycin and Torin1-dependent MDC induction was reduced in low amino acid medium, blocked in amino acid free medium, and restored with casamino acids (Figures 4E and S4G). These results demonstrate that high intracellular amino acid concentrations trigger MDC formation through an mTOR-independent mechanism, and that inhibiting mTOR induces MDCs by elevating intracellular amino acids. In addition to the mTOR pathway, we also tested whether genetic alteration of other common amino acid sensing pathways in yeast impacted MDC formation. We found that deletion of *GCN2*, *GLN3*, *GPA2*, *GPR1*, and *SSY1*, which dismantle a variety of nutrient signaling system in yeast, had no impact on MDC formation in response to concA, CHX, or rapamycin (Figure 4F). Thus, amino acid surplus triggers MDC formation independently of common nutrient sensing pathways.

### MDCs regulate the abundance of the carrier receptor Tom70 on mitochondria

Through microscopy-based screening, we previously found that the cargo localized to MDCs included Tom70, a mitochondrial OM receptor that facilitates import of the SLC25 nutrient transporter family into mitochondria (Figure S5E) (Sollner et al., 1990), members of the inner membrane-localized SLC25 family themselves, as well as a few other OM proteins that are known to be Tom70 clients (Hughes et al., 2016). The SLC25 protein family consists of over 30 members in yeast and 50-plus members in mammals, and these multi-pass IM proteins are responsible for the exchange of nearly all metabolites across the mitochondrial IM (Palmieri and Monne, 2016). Because MDCs are nutrient responsive, we wondered whether these structures functioned to modulate the Tom70 pathway in mitochondria by reducing Tom70 levels on the mitochondrial surface upon nutrient surplus. Consistent with this idea, we found that after rapamycin treatment, Tom70 was nearly 6-fold enriched in the MDC compared to the mitochondrial tubule (Figure 5A-B). This enrichment led to a ∼50% reduction in the abundance of Tom70 in the mitochondrial tubule after two hours (Figure 5C). Similar decreases in mitochondrial tubule abundance of Tom70 were observed across all known MDC inducers (Figure S5A). By contrast, mitochondrial levels of Tim50 and Ilv2, IM and matrix proteins which do not associate with MDCs, were not reduced upon treatment with MDC inducers (Figures S5B-D). Importantly, in an accompanying manuscript, we found that the conserved mitochondrial localized GTPase Gem1 (Frederick et al., 2004) is required for MDC formation (English et al., 2020). Ablation of MDC formation by deleting *GEM1* prevented removal of Tom70 from mitochondria in response to MDC activating treatments (Figures 5C and S5A), indicating that MDCs diminish the abundance of Tom70 in mitochondria under elevated amino acids conditions.

**Figure 5.**
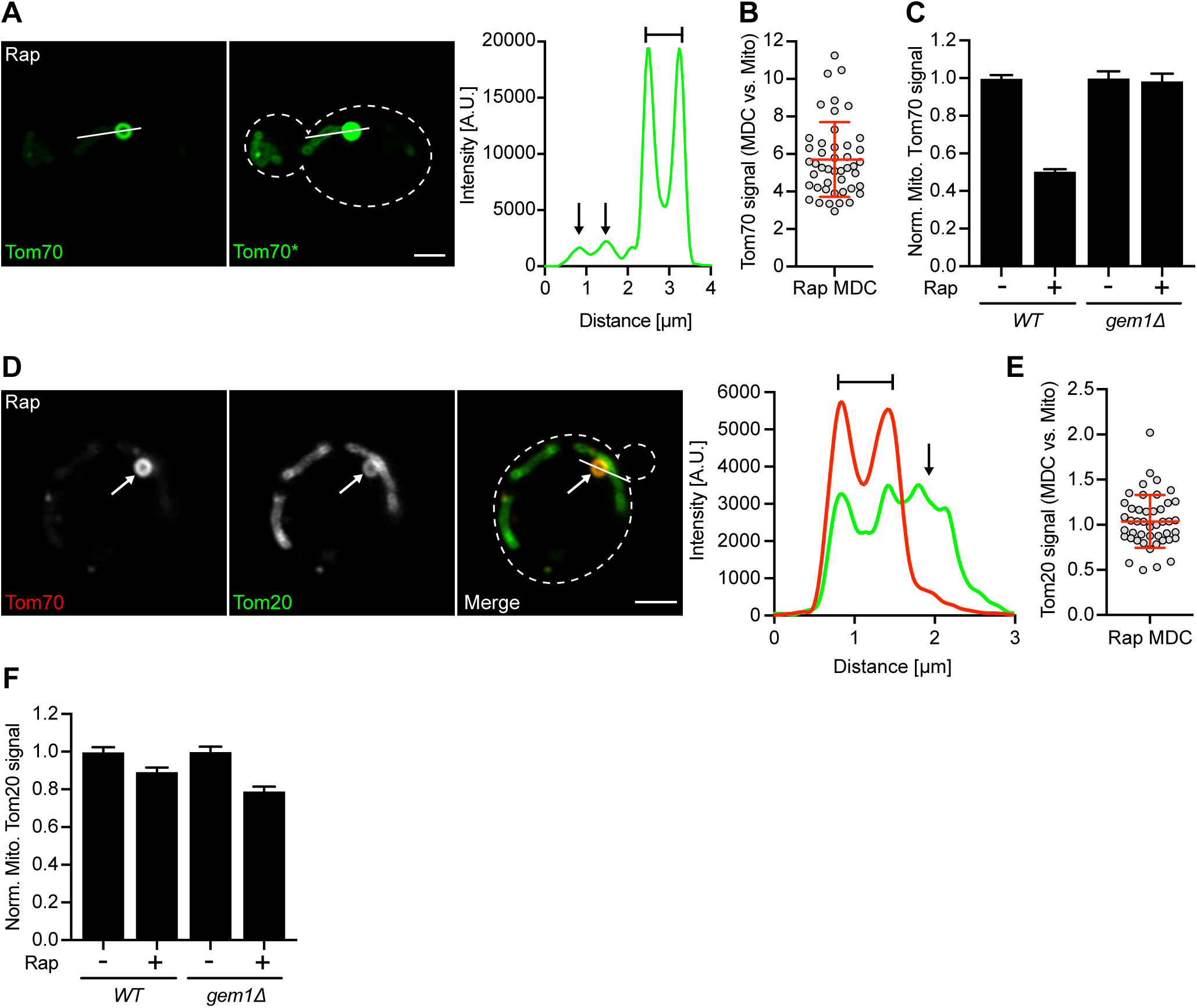
MDCs Selectively Sequester the Carrier Receptor Tom70 from Mitochondria. (A) Super-resolution images (left) and line-scan analysis (right) of rapamycin (Rap)-induced MDC formation in yeast cells expressing Tom70-GFP. Right image (Tom70*) shows where the fluorescence intensity has been increased post imaging to visualize the mitochondrial tubule. White line marks fluorescence intensity profile position. Left line-scan Y axis corresponds to the Tom70-GFP fluorescence intensity. Black arrow marks mitochondrial tubule. Bracket marks MDC. Scale bar = 2 µm. (B) Scatter plot showing the normalized Tom70-GFP fluorescence intensity in Rap-induced MDCs compared to the adjacent mitochondrial tubule. Error bars show mean ± SD of *n* = 45 cells from three per replicates with *n* = 15 cells per replicate. (C) Normalized mitochondrial Tom70-GFP fluorescence in *wild-type (WT)* and *gem1Δ* cells treated with Rap compared to DMSO. Error bars show mean ± SE of three replicates with *n* = 15 cells per replicate. (D) Super-resolution images (left) and line-scan analysis (right) of Rap-induced MDC formation in yeast cells expressing Tom20-GFP and Tom70-mCherry. White arrow marks MDC. White line marks fluorescence intensity profile position. Line-scan Y axis corresponds to Tom20-GFP and Tom70-mCherry fluorescence intensity. Black arrow marks mitochondrial tubule. Bracket marks MDC. Scale bar = 2 µm. (E) Scatter plot showing the normalized Tom20-GFP fluorescence intensity in Rap-induced MDCs compared to the adjacent mitochondrial tubule. Error bars show mean ± SD of *n* = 45 cells from three per replicates with *n* = 15 cells per replicate. (F) Normalized mitochondrial Tom20-GFP fluorescence in *wild-type (WT)* and *gem1Δ* cells treated with Rap or DMSO. Error bars show mean ± SE of three replicates with *n* = 15 cells per replicate. See also Figure S5.

In addition to the carrier import receptor Tom70, mitochondria also harbor another OM import receptor, Tom20, which binds and facilitates import of matrix and IM localized proteins that contain canonical N-terminal mitochondrial targeting sequences (MTSs) (Figure S5E) (Abe et al., 2000; Moczko et al., 1994; Moczko et al., 1993). In contrast to Tom70 and its clients, we previously found via microscopy screening that Tom20 and all MTS containing proteins were excluded from MDCs (Hughes et al., 2016). Using super resolution microscopy, we confirmed this observation here. We found that while Tom20 is present at low levels in MDCs, it is not enriched above the signal in the mitochondrial tubule, and is not depleted from the mitochondrial tubule under any MDC inducing condition (Figures 5D-F and S5F). These results indicate that MDCs specifically target the Tom70 pathway in response to elevated amino acids, while leaving the Tom20 pathway and its clients unchanged.

### MDCs are regulated by mitochondrial nutrient transporter abundance

Given that MDCs sequester machinery and clients of the Tom70 carrier pathway in response to elevated amino acids (Hughes et al., 2016), we reasoned that modulation of SLC25 transporter levels may in turn regulate MDC formation. Consistent with this idea, overexpression of Tom70, which provides more binding sites for carriers on the mitochondrial surface, caused constitutive MDC formation in cells grown in high amino acid medium, and stimulated MDC formation in concA-treated cells grown in low amino acid medium, where MDC formation is normally not apparent (Figure 6A). Moreover, deletion of *TOM70* (Hines et al., 1990; Steger et al., 1990) alone or in combination with its paralog *TOM71* (Schlossmann et al., 1996), which prevents localization of carriers to mitochondria (Ryan et al., 1999), severely reduced MDC formation in response to all known MDC inducers (Figure 6B). By contrast, MDC formation was largely unaffected in cells overexpressing or deficient for *TOM20* (Figures 6C-D), again highlighting the specificity of the MDC pathway for Tom70.

**Figure 6.**
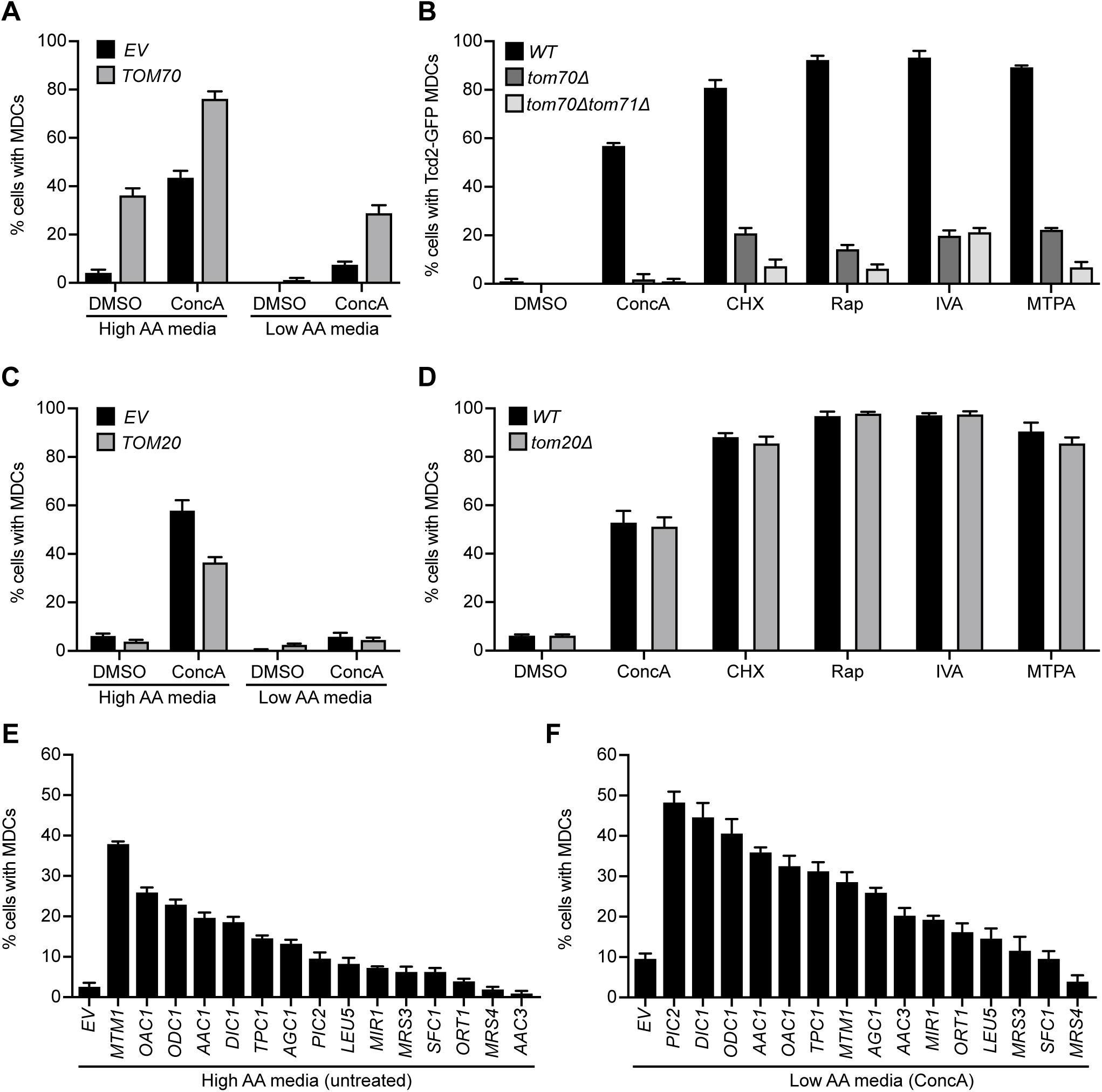
Mitochondrial Carrier Levels Control MDC Formation. (A) Quantification of concanamycin A (ConcA)-induced MDC formation in *TOM70* overexpressing or *empty vector* (*EV*) control cells in high and low amino acid media. Error bars show mean ± SE of three replicates with *n* = 100 cells per replicate. (B) Quantification of ConcA, cycloheximide (CHX), rapamycin (Rap), isovaleraldehyde (IVA) and 3-methylthiopropanal (MTPA)-induced MDC formation in *wild-type (WT)*, *tom70Δ* and *tom70Δ tom71Δ* cells. Error bars show mean ± SE of three replicates with *n* = 100 cells per replicate. (C) Quantification of ConcA-induced MDC formation in *TOM20* overexpressing or *EV* control cells in high and low amino acid media. Error bars show mean ± SE of three replicates with *n* = 100 cells per replicate. (D) Quantification of ConcA, CHX, Rap, IVA and MTPA-induced MDC formation in *WT* and *tom20Δ* cells. Error bars show mean ± SE of three replicates with *n* = 100 cells per replicate. (E) Quantification of MDC formation in cells expressing the indicated mitochondrial carrier in high amino acid media in absence of drug treatment. Error bars show mean ± SE of three replicates with *n* = 100 cells per replicate. (E) Quantification of MDC formation in cells expressing the indicated mitochondrial carrier in low amino acid media in presence of ConcA. Error bars show mean ± SE of three replicates with *n* = 100 cells per replicate. See also Figure S6.

To more directly analyze the impact of carrier abundance on the MDC pathway, we overexpressed 28 individual members of the SLC25 family in wild-type cells, and screened for carriers that impacted the magnitude of MDC formation (Figures S6A-B). We identified numerous SLC25 carriers that constitutively activated MDC formation when overexpressed in cells grown in high amino acid medium, and stimulated MDC formation in concA-treated cells grown in low amino acid medium (Figures 6E-F and S6A-B). Interestingly, several of these carriers play direct or indirect roles in amino acid metabolism. By contrast, overexpression of the non-MDC IM client Tim50 did not activate MDC formation (Figure S6C). Collectively, these results suggest that the MDC pathway selectively targets the Tom70 carrier pathway in response to amino acid alterations, and that modulating mitochondrial carrier levels via direct carrier overexpression, or by regulating the abundance of the carrier receptor Tom70 governs the extent of MDC formation within cells.

### MDCs act in parallel with the lysosome and the MVB pathway to protect cells from amino acid overload

Finally, we wanted to determine whether MDCs play a role in regulating cellular health under conditions of elevated amino acid stress. We first tested whether deletion of *GEM1*, which is required for MDC formation (English et al., 2020), impacted cell growth under MDC inducing conditions. Serial-dilution growth assays demonstrated mild to no growth defects in *gem1*Δ strains when grown in the presence of concA combined with leucine or methionine addition—conditions that activate the MDC pathway (Figure 7A). Since *gem1*Δ strains did not exhibit robust growth phenotypes under MDC inducing conditions, we considered the existence of redundant pathways that protect cells from amino acid stress in absence of MDC formation. A strong candidate for such a system is the endosomal sorting complexes required for transport (ESCRT)-dependent MVB pathway. This pathway post-translationally regulates levels of nutrient transporters at the plasma membrane (PM) via ubiquitin-mediated receptor internalization and destruction in the lysosome, and mutants in this system exhibit elevated amino acid uptake from the environment (Katzmann et al., 2002; Rubio-Texeira and Kaiser, 2006).

**Figure 7.**
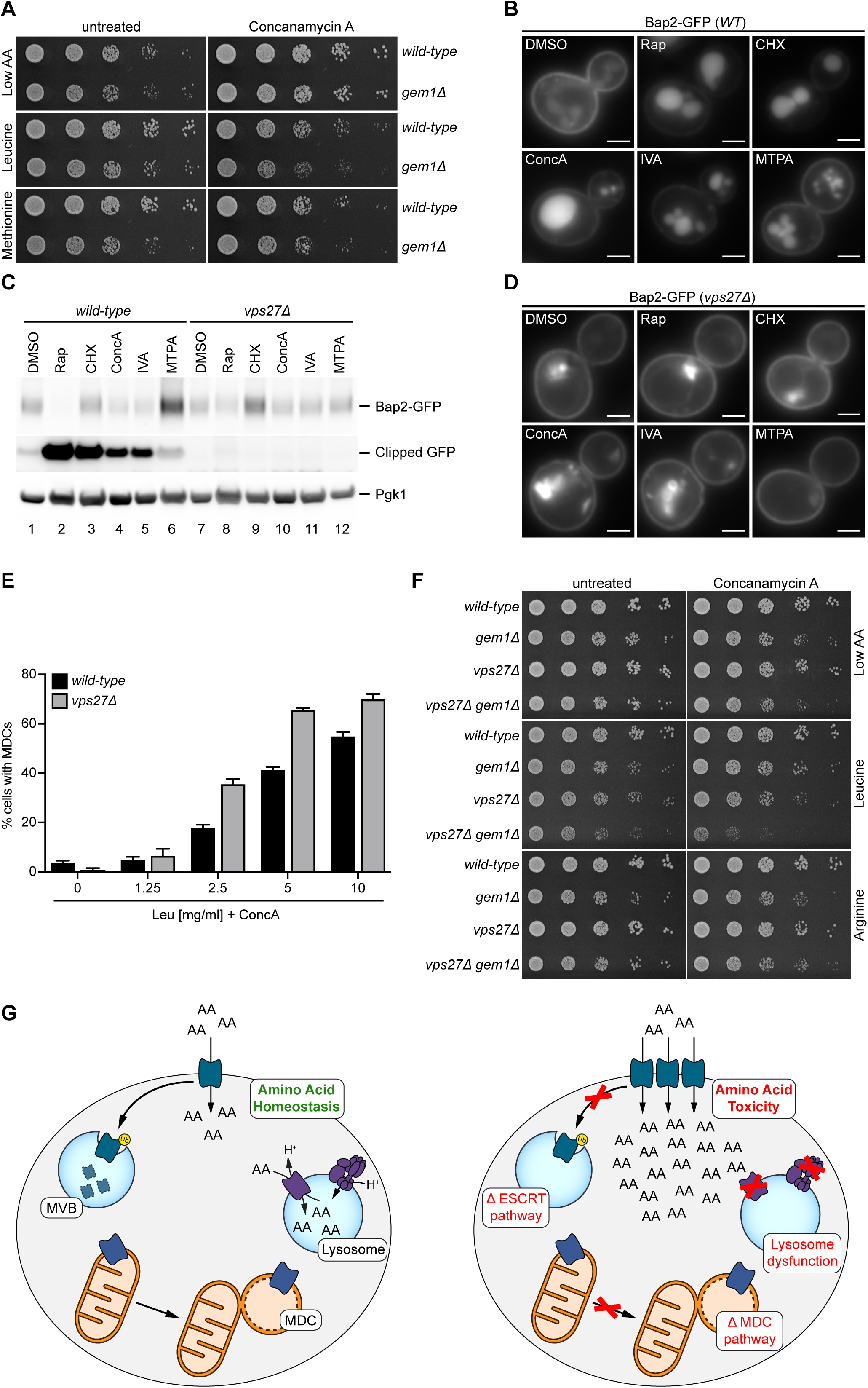
MDCs Cooperate with MVBs and Lysosomes to Protect Cells from Toxic Levels of Amino Acids. (A) Growth of *wild-*type and *gem1Δ* cells in presence and absence of concanamycin A (ConcA) on low amino acid media supplemented with 10mg/ml leucine or methionine. (B) Widefield images showing internalization of Bap2-GFP upon rapamycin (Rap), cycloheximide (CHX), ConcA, isovaleraldehyde (IVA) or 3-methylthiopropanal (MTPA) treatment. Scale bar = 2 µm. (C) Western blot analysis of Bap2-GFP clipping upon treatment with Rap, CHX, ConcA, IVA and MTPA in *wild-type* and *vps27Δ* cells. Pgk1 – loading control. (D) Widefield images showing stabilization of Bap2-GFP on the plasma membrane in Rap, CHX, ConcA, IVA or MTPA treated *vps27Δ* cells. Bright structures resemble typical class E compartments found in ESCRT mutants. Scale bar = 2 µm. (E) Quantification of ConcA-induced MDC formation in low amino acid media supplemented with the indicated amount of leucine in *wild-type (WT)* and *vps27Δ* cells. Error bars show mean ± SE of three replicates with *n* = 100 cells per replicate. (F) Growth of *wild-type*, *gem1Δ, vps27Δ* and *gem1Δ vps27Δ* strains in presence and absence of ConcA on low amino acid media supplemented with 10mg/ml leucine or arginine. (G) Yeast cells utilize multiple systems, including lysosomal amino acid compartmentation, ESCRT-dependent control of plasma-membrane localized amino acid transporter via the MVB pathway and the MDC pathway to counteract toxic levels of amino acids. Simultaneous defects in all three systems render cells sensitive to toxic levels of amino acids. See also Figure S7.

Prior studies showed that the ESCRT/MVB system downregulates PM transporters in response to rapamycin and CHX—two conditions that also activate MDC formation (Hatakeyama and De Virgilio, 2019; Lin et al., 2008; Nikko and Pelham, 2009). Utilizing the fluorescently labelled branched-chain amino acid permease Bap2-GFP (Grauslund et al., 1995), we confirmed via microscopy and western blot that treatment with rapamycin or CHX triggered a reduction in Bap2-GFP levels at the PM, and a corresponding increase in Bap2-GFP processing in the lysosome—the latter indicated by an accumulation of free GFP that is proteolytically released from Bap2-GFP in the lysosome (Figures 7B-C). Remarkably, Bap2 internalization and processing also occurred in other conditions that activate MDCs, including concA, or treatment with the leucine and methionine derived aldehydes IVA and MTPA (Figures 7B-C). We confirmed that concA-induced Bap2 internalization and degradation was prevented in cells lacking components of the ESCRT machinery (Hurley and Emr, 2006), including *VPS27* and *DID4* (Figures 7C-D and S7A-B). Thus, under conditions of amino acid excess, cells simultaneously activate MDC formation and reduce levels of PM nutrient transporters via the ESCRT-dependent MVB pathway.

Based on these observations, we reasoned that the ESCRT-dependent MVB pathway and MDCs may act in parallel to protect cells from amino acid stress. Consistent with this idea, MDC formation was elevated in cells lacking the ESCRT components *VPS27* and *DID4* (Figures 7E and S7C). Moreover, while cells lacking either *GEM1* or components of the ESCRT machinery exhibited mild to no growth defects in the presence of high leucine or methionine, simultaneous loss of both the MDC and the ESCRT-dependent MVB pathways was detrimental under high amino acid conditions, and severely inhibited growth when combined with loss of lysosome function (Figures 7F and S7D-E). Importantly, growth inhibition only occurred in response to the MDC inducing amino acids leucine and methionine, and was not caused by arginine or proline, two amino acids that do not activate MDC formation (Figures 7F and S7D-E). Altogether, these results suggest that MDCs act in parallel with lysosomes and the ESCRT-dependent MVB pathway to ensure cellular survival under conditions of amino acid stress.

## DISCUSSION

Here, we demonstrate that a previously identified foci associated with mitochondria, the MDC (Hughes et al., 2016), is a dynamic, lumen-containing organelle that is generated from mitochondria in response to intracellular amino acid overload. Upon formation, MDCs selectively sequester the machinery and cargo of the Tom70 metabolite carrier import pathway away from the rest of the mitochondrial network. Based on our data, we speculate that MDCs function to protect cells from amino acid stress, potentially by acting as part of a broad cellular network that coordinately regulates intracellular nutrient distribution. The three nodes of this network, as depicted in Figure 7G, are the ESCRT-dependent MVB pathway that controls metabolite transporters at the PM (Katzmann et al., 2002), the lysosome, which sequesters and stores amino acids in its lumen (Klionsky et al., 1990; Wiemken and Dürr, 1974), and the MDC, which regulates the metabolite carrier pathway on mitochondria. Consistent with the idea that these systems act in parallel to preserve intracellular amino acid homeostasis, combined loss of the MDC pathway, the ESCRT-dependent MVB system, and lysosome amino acid storage leads to cellular toxicity that is amino acid dependent. These results highlight the importance of maintaining proper amino acid homeostasis in cells, and suggest cells contain dedicated systems to mitigate toxicity associated with improper levels of amino acids.

The discovery of a new nutrient-sensitive compartment raises many important questions for future exploration. Central among them is how MDCs prevent toxicity from excess BCAAs and methionine? Our data indicate that MDCs negatively regulate the levels of Tom70, an OMM protein required for import of nutrient carriers into mitochondria, and that the MDC pathway is in turn sensitive to carrier expression levels. Thus, we speculate that much like the MVB-dependent ESCRT pathway that regulates transporter levels on the PM, the MDC in turn modulates metabolite carrier levels on the mitochondria in response to changes in cellular nutrient supply. A key difference between the PM and mitochondrial systems is the fact that mitochondrial nutrient carriers are localized on the mitochondrial IM, which is not accessible from the cytoplasm. Thus, sequestering the import receptor on the mitochondrial surface, Tom70, may enable cells to indirectly control mitochondrial nutrient transporters. Additional studies are required to solidify whether the MDC incorporates both the mitochondrial OM and IM, and whether carrier proteins localized to the MDC are enriched from the IM.

Regardless of how the MDC protects cells from amino acid stress, our data strongly suggests a role for mitochondria in mediating the toxic effects of excess BCAAs and methionine. We recently discovered that defects in cysteine homeostasis cause cellular toxicity by altering iron availability and mitochondrial respiration (Hughes et al., 2020). However, MDCs do not respond to cysteine or iron perturbations, indicating that the mechanism by which excess BCAAs and methionine become toxic to cells are distinct from cysteine’s impact on mitochondrial respiration. Interestingly, elevated BCAAs have been linked to the development of metabolic disorders including Type II diabetes (Knebel et al., 2016; Newgard et al., 2009; Wang et al., 2011; Xu et al., 2013). Current speculation suggests that elevated BCAAs or their catabolites may alter insulin production by impairing mitochondrial function (Lynch and Adams, 2014). However, the mechanism by which this occurs remains unknown. Deciphering the mode of action of BCAA and methionine toxicity is an important area of future research, as well as understanding how MDCs function to protect cells from the toxic effects of these amino acids.

We show that MDCs are responsive to elevated levels of intracellular amino acids, with BCAAs and downstream catabolites being the most potent MDC inducers. It remains to be determined how signals arising from elevated amino acids are relayed to the MDC formation machinery. We anticipate that clues to this question may be garnered from studies on other nutrient responsive systems, including the autophagosome and MVB pathways. Each of these systems contains machinery that integrate nutrient cues into remodeling of cellular membranes. In contrast to MDCs and the MVB pathway, autophagosomes are generated in response to nutrient deprivation, and nutrient cues are relayed to the autophagosome machinery via well-characterized nutrient sensing systems such as mTOR (Saxton and Sabatini, 2017; Yin et al., 2016). At this point, it appears that MDCs are activated in the exact opposite scenario as autophagosomes—under nutrient replete conditions. Interestingly, all treatments that stimulate MDC production also trigger downregulation of nutrient transporters via the MVB pathway (Figures 7B-C). Thus, it seems likely that these two pathways utilize similar nutrient sensing systems—the identify of which is unknown at this time. Moving forward, it will be important to determine how the MDC and MVB systems act coordinately with lysosomes and protein translation machinery to regulate cellular amino acid homoeostasis, and how deficiencies in these systems impact cellular health.

## ACKNOWLEDGEMENTS

We thank members of the A.L.H. and J.M.S laboratories for discussion and manuscript comments and L. VanderMeer (Utah) and P. Guo (Utah) for technical assistance. Metabolomics analysis was performed at the University of Utah Metabolomics Core directed by J. Cox and supported by National Institutes of Health (NIH) grants 1S10OD016232-01, 1S10OD021505-01 and 1U54DK110858-01. Research was supported by NIH grants AG043095, GM119694, AG061376, and AG055648 (A.L.H.), NIH T32GM007464 (A.M.E.), NIH GM53466 and GM84970 (J.M.S.), AHA 18PRE33960427 (M.H.S.) and the Howard Hughes Medical Institute (J.M.S.). A.L.H. was further supported by an American Federation for Aging Research Junior Research Grant, United Mitochondrial Disease Foundation Early Career Research Grant, Searle Scholars Award, and Glenn Foundation for Medical Research Award.

## AUTHOR CONTRIBUTIONS

Conceptualization, M.H.S., A.M.E., A.L.H., and J.M.S.; Methodology, M.H.S., A.M.E., A.L.H., and J.M.S.; Formal Analysis, M.H.S., A.M.E., and T.J.C.; Investigation, M.H.S., A.M.E., and T.J.C.; Writing - Original Draft, A.L.H., A.M.E., and M.H.S.; Writing - Reviewing and Editing, A.L.H. and J.M.S.; Visualization, M.H.S. and A.M.E.; Supervision, A.L.H. and J.M.S.; Funding Acquisition, A.L.H., J.M.S., A.M.E., and M.H.S.

---

**Figure S1.**
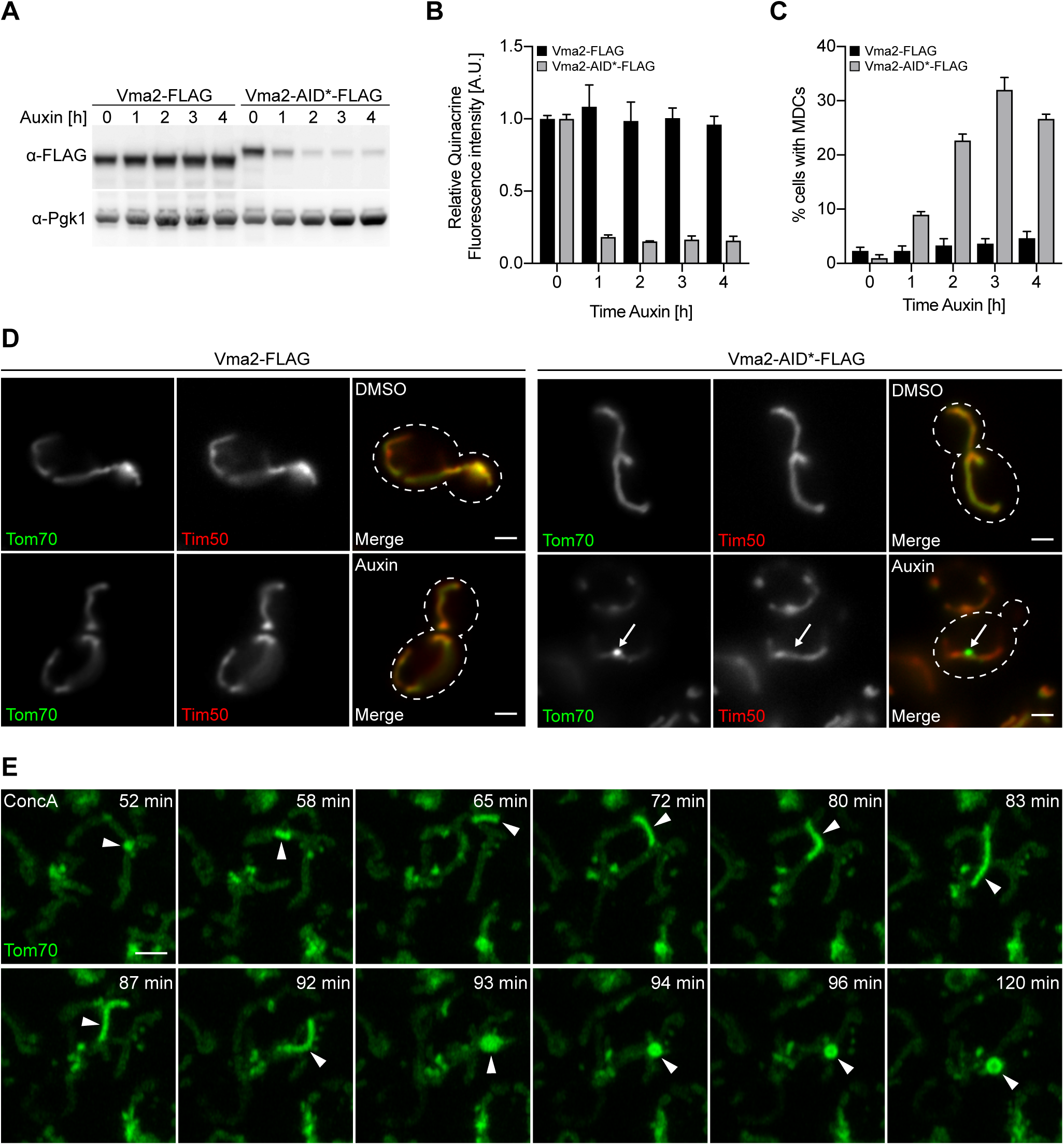
V-ATPase Inhibition Induces Dynamic MDC Formation, Related to Figure 1. (A) Western blot showing time course of auxin-induced Vma2-AID*-FLAG depletion. (B) Time course of auxin-induced inhibition of vacuole acidification due to Vma2-AID*-FLAG depletion assayed by staining with pH-dependent dye quinacrine. Error bars show mean ± SE of three replicates, *n* = 30 cells per replicate. (C) Quantification of Vma2-AID*-FLAG-depletion induced MDC formation over time. Error bars show mean ± SE of three replicates with *n* = 100 cells per replicate. (D) Widefield images of auxin-induced MDC formation in the indicated yeast strains expressing Tom70-GFP and Tim50-mCherry. White arrow marks MDC. Scale bar = 2 µm. (E) Time-lapse images of ConcA-induced MDC formation in yeast cells expressing Tom70-GFP. Images were acquired over 120 minutes (min). Arrowhead marks MDC. Scale bar = 2 µm. See also Video S2.

**Figure S2.**
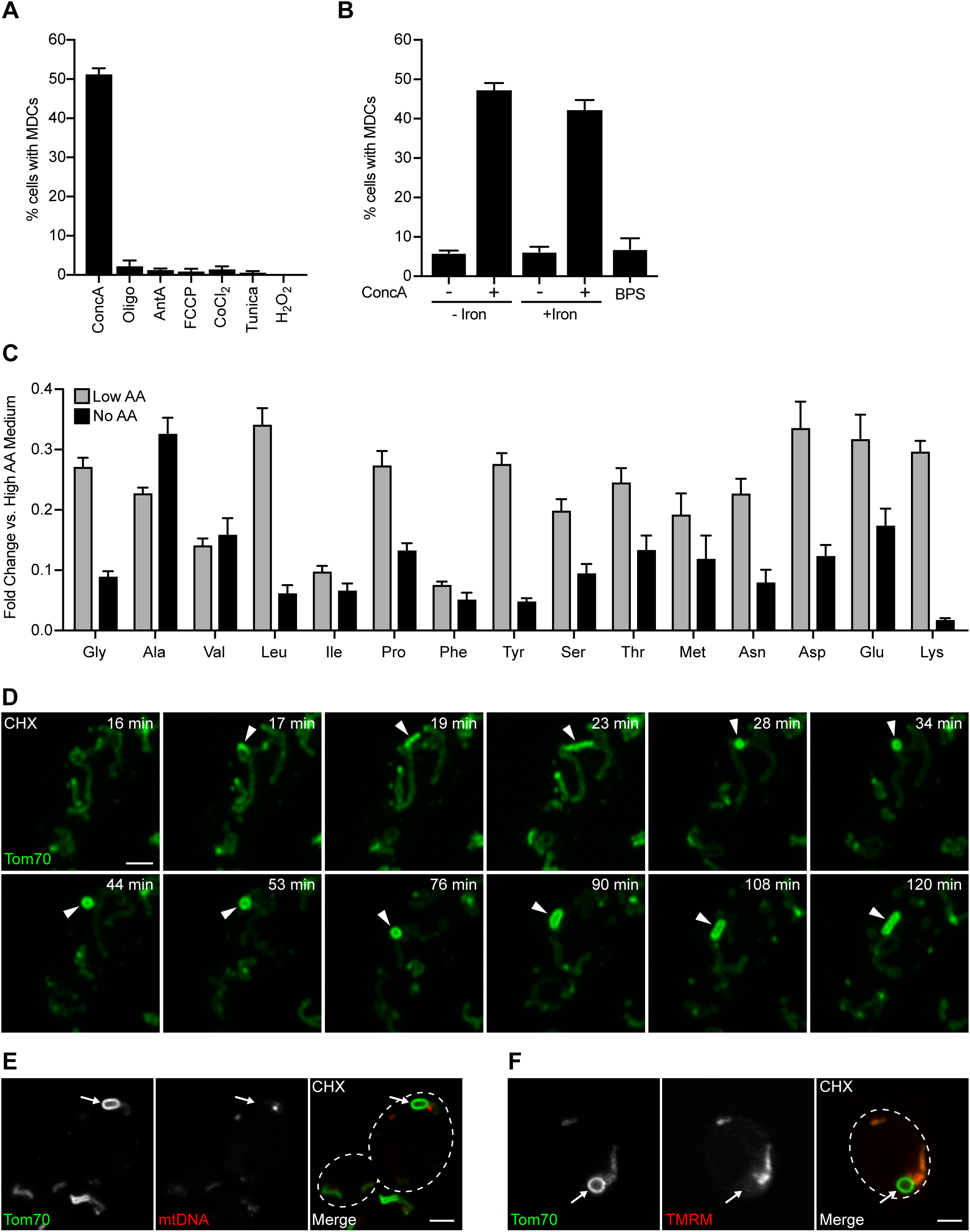
Additional Characteristics of MDCs, Related to Figure 2. (A) Quantification of MDC formation in yeast cells treated with concanamycin A (ConcA), oligomycin (Oligo), antimycin A (AntA), carbonyl cyanide 4- (trifluoromethoxy)phenylhydrazone (FCCP), CoCl_2_, tunicamycin (Tunica) or H_2_O_2_. Error bars show mean ± SE of three replicates with *n* = 100 cells per replicate (B) Quantification of MDC formation in yeast cells treated with ConcA or bathophenanthroline disulfonate (BPS) in high amino acid media in presence or absence of exogenous iron. Error bars show mean ± SE of three replicates with *n* = 100 cells per replicate. (C) Relative amino acid levels in yeast cells grown in media containing low or no amino acids compared to high amino acid medium assayed by GC-MS. Error bars show mean ± SE of three replicates. (D) Time-lapse images of CHX-induced MDC formation in yeast cells expressing Tom70-GFP. Images were acquired over 120 minutes (min). Arrowhead marks MDC. Scale bar = 2 µm. See also Video S3. (E) Super-resolution images of cycloheximide (CHX) treated yeast cells expressing Tom70-GFP stained with DAPI to label mitochondrial DNA. White arrow marks MDC. Scale bar = 2 µm. (F) Super-resolution images of CHX treated yeast cells expressing Tom70-GFP stained with fluorescent membrane potential indicator TMRM. White arrow marks MDC. Scale bar = 2 µm.

**Figure S3.**
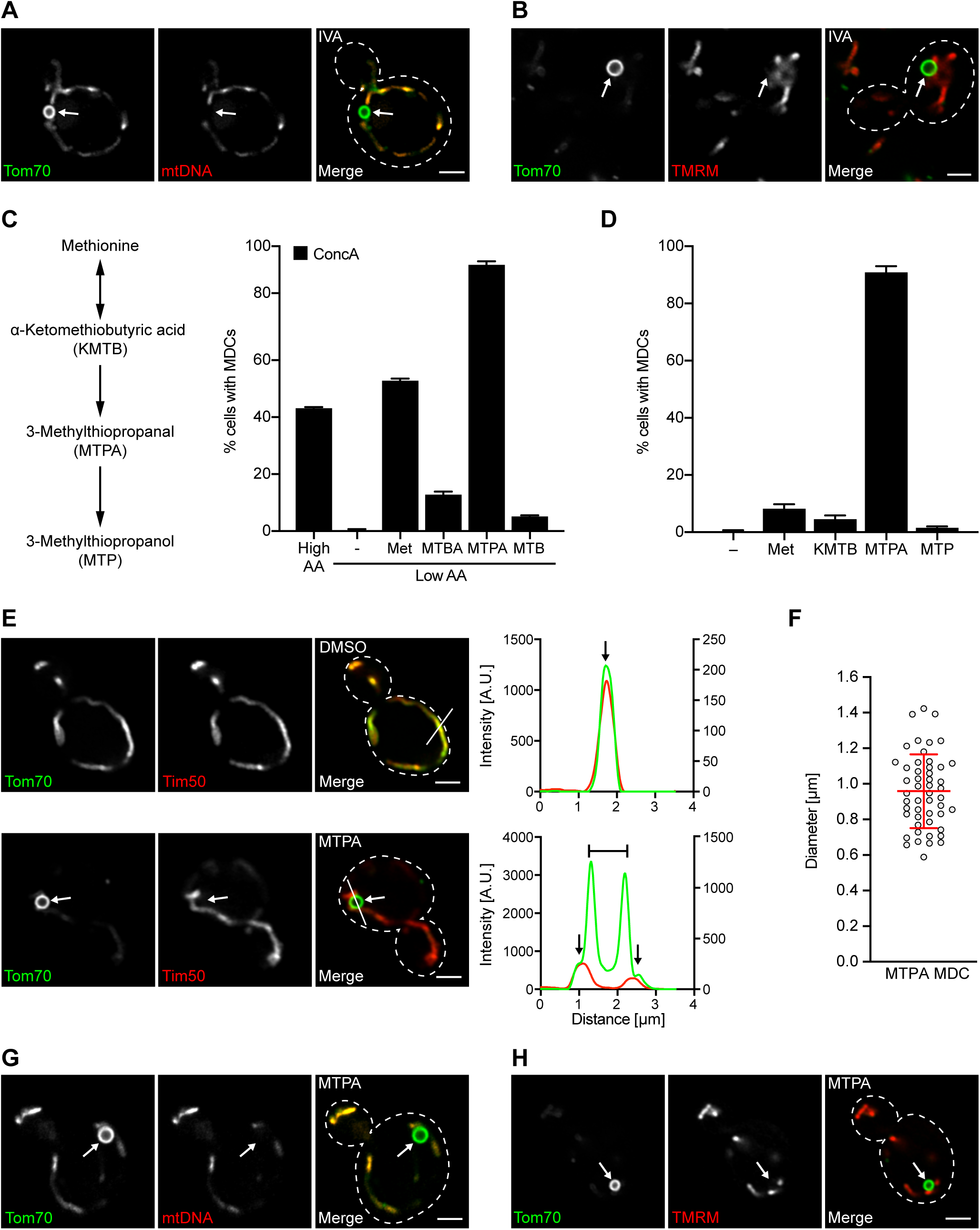
Leucine and Methionine Derivatives Activate MDC Formation, Related to Figure 3. (A) Super-resolution images of isovaleraldehyde (IVA) treated yeast cells expressing Tom70-GFP, stained with DAPI to label mitochondrial DNA. White arrow marks MDC. Scale bar = 2 µm. (B) Super-resolution images of IVA treated yeast cells expressing Tom70-GFP stained with fluorescent membrane potential indicator TMRM. White arrow marks MDC. Scale bar = 2 µm. (C) Quantification of concanamycin A (ConcA)-induced MDC formation in low amino acid medium supplemented with methionine (Met) or its catabolites α-ketomethiobutyric acid (KMTB), 3-methylthiopropanal (MTPA) or 3-methylthiopropanol (MTP). Error bars show mean ± SE of three replicates with *n* = 100 cells per replicate. (D) Quantification of MDC formation in high amino acid medium supplemented with Met, KMTB, MTPA or MTP in the absence of drug treatment. Error bars show mean ± SE of three replicates with *n* = 100 cells per replicate. (E) Super-resolution images (left) and line-scan analysis (right) of MTPA-induced MDC formation in yeast cells expressing Tom70-GFP and Tim50-mCherry. White arrow marks MDC. White line marks fluorescence intensity profile position. Left and right line-scan Y axis correspond to Tom70-GFP and Tim50-mCherry fluorescence intensity, respectively. Black arrow marks mitochondrial tubule. Bracket marks MDC. Scale bar = 2 µm. (F) Scatter plot showing the diameter of MTPA-induced MDCs. Error bars show mean (0.9587 µm) ± SD of *n* = 50 MDCs. (G) Super-resolution images of MTPA-induced MDC formation in yeast cells expressing Tom70-GFP, stained with DAPI to visualize mitochondrial DNA. White arrow marks MDC. Scale bar = 2 µm. (H) Super-resolution images of MTPA-induced MDC formation in yeast cells expressing Tom70-GFP, stained with fluorescent membrane potential indicator TMRM. White arrow marks MDC. Scale bar = 2 µm.

**Figure S4.**
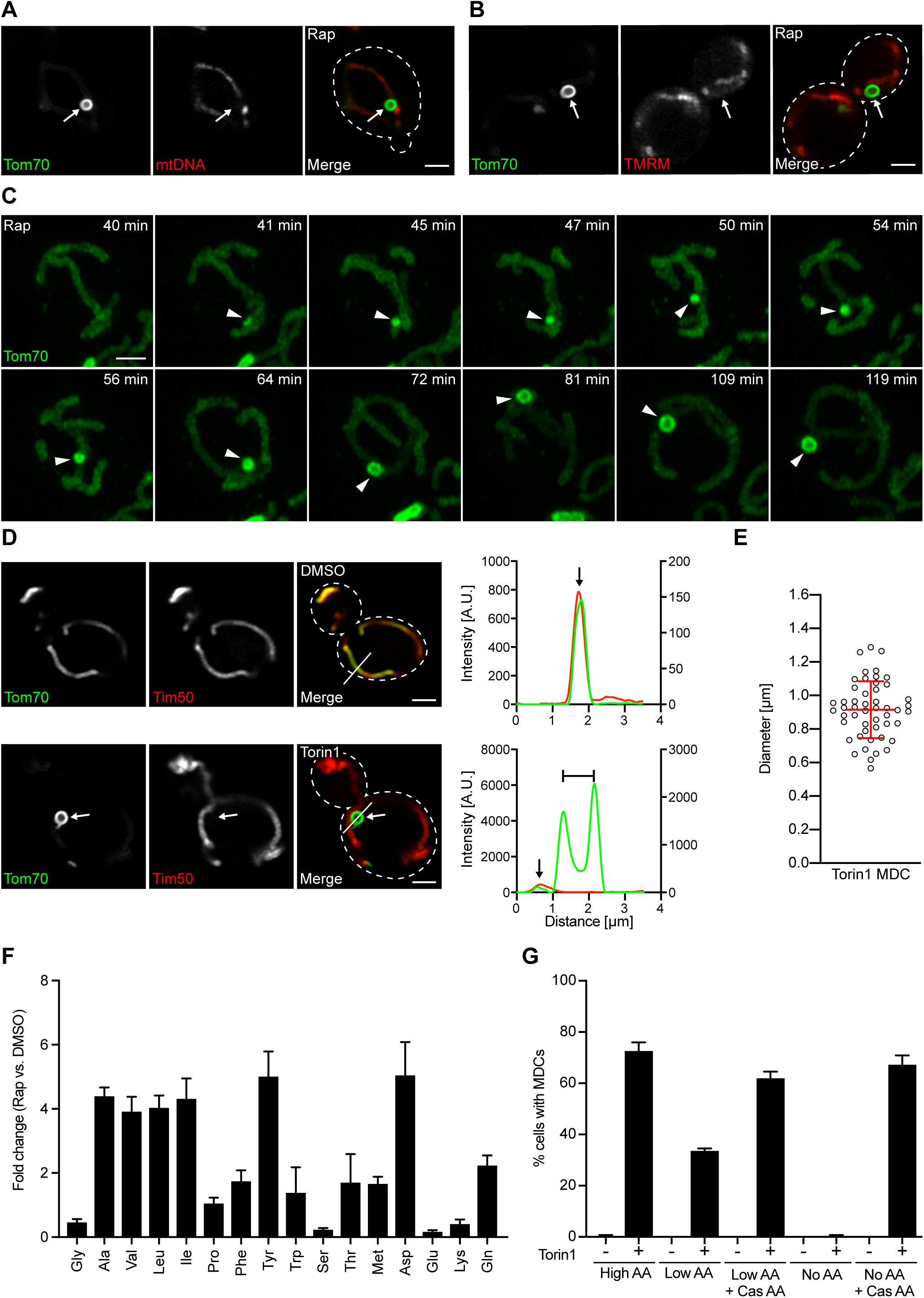
Rapamycin and Torin1 Treatment Activate MDC Formation, Related to Figure 4. (A) Super-resolution images of Rap-induced MDC formation in yeast cells expressing Tom70-GFP, stained with DAPI to visualize mitochondrial DNA. White arrow marks MDC. Scale bar = 2 µm. (B) Super-resolution images of Rap-induced MDC formation in yeast cells expressing Tom70-GFP, stained with fluorescent membrane potential indicator TMRM. White arrow marks MDC. Scale bar = 2 µm. (C) Time-lapse images showing Rap-induced MDC formation in yeast cells expressing Tom70-GFP. Images were acquired over 119 minutes (min). Arrowhead marks MDC. Scale bar = 2 µm. See also Video S4. (D) Super-resolution images (left) and line-scan analysis (right) of Torin1-induced MDC formation in yeast cells expressing Tom70-GFP and Tim50-mCherry. White arrow marks MDC. White line marks fluorescence intensity profile position. Left and right line-scan Y axis correspond to Tom70-GFP and Tim50-mCherry fluorescence intensity, respectively. Black arrow marks mitochondrial tubule. Bracket marks MDC. Scale bar = 2 µm. (E) Scatter plot showing the diameter of Torin1-induced MDCs. Error bars show mean (0.9156 µm) ± SD of *n* = 50 MDCs. (F) Relative amino acid levels in Rap treated yeast cells grown in medium containing high levels amino acids assayed by GC-MS. Error bars show mean ± SE of three replicates. (G) Quantification of Torin1-induced MDC formation in media containing high, low or no amino acids (AA). Cas AA indicates where bulk amino acids were added back to media in the form of hydrolyzed casein. Error bars show mean ± SE of three replicates with *n* = 100 cells per replicate.

**Figure S5.**
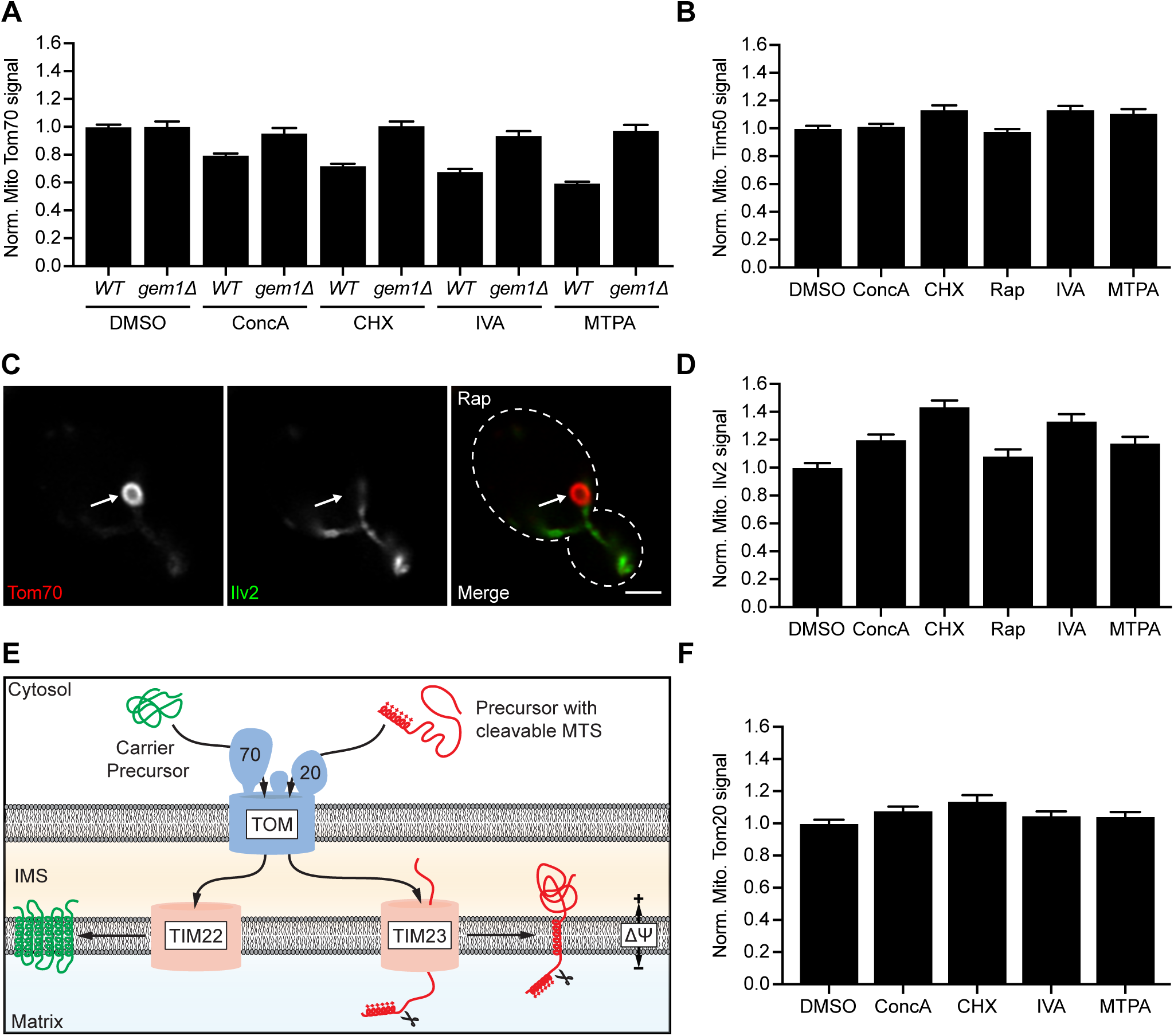
MDCs Selectively Remove Tom70, Related to Figure 5. (A) Normalized mitochondrial Tom70-GFP fluorescence in *wild-type (WT)* and *gem1Δ* cells treated with concanamycin A (ConcA), cycloheximide (CHX), isovaleraldehyde (IVA) or 3-methylthiopropanal (MTPA)-compared to DMSO. Error bars show mean ± SE of three replicates with *n* = 15 cells per replicate. This graph shows additional data generated in the experiments used for Figure 5C. The DMSO control is duplicated from Figure 5C for comparison purposes. (B) Normalized mitochondrial Tim50-GFP fluorescence in cells treated with ConcA, CHX, rapamycin (Rap), IVA or MTPA compared to DMSO. Error bars show mean ± SE of three replicates with *n* = 15 cells per replicate. (C) Super-resolution images of Rap-induced MDC formation in yeast cells expressing Ilv2-GFP and Tom70-mCherry. White arrow marks MDC. Scale bar = 2 µm. (D) Normalized mitochondrial Ilv2-GFP fluorescence in cells treated with ConcA, CHX, Rap, IVA or MTPA compared to DMSO. Error bars show mean ± SE of three replicates with *n* = 15 cells per replicate. (E) Model of mitochondrial protein import. Tom70 promotes carrier biogenesis, whereas Tom20 is required for protein import of pre-sequence containing proteins into mitochondria. (F) Normalized mitochondrial Tom20-GFP fluorescence in cells treated with ConcA, CHX, IVA or MTPA compared to DMSO. Error bars show mean ± SE of three replicates with *n* = 15 cells per replicate. This graph shows additional data generated in the experiments used for Figure 5F. The DMSO control is duplicated from Figure 5F for comparison purposes.

**Figure S6.**
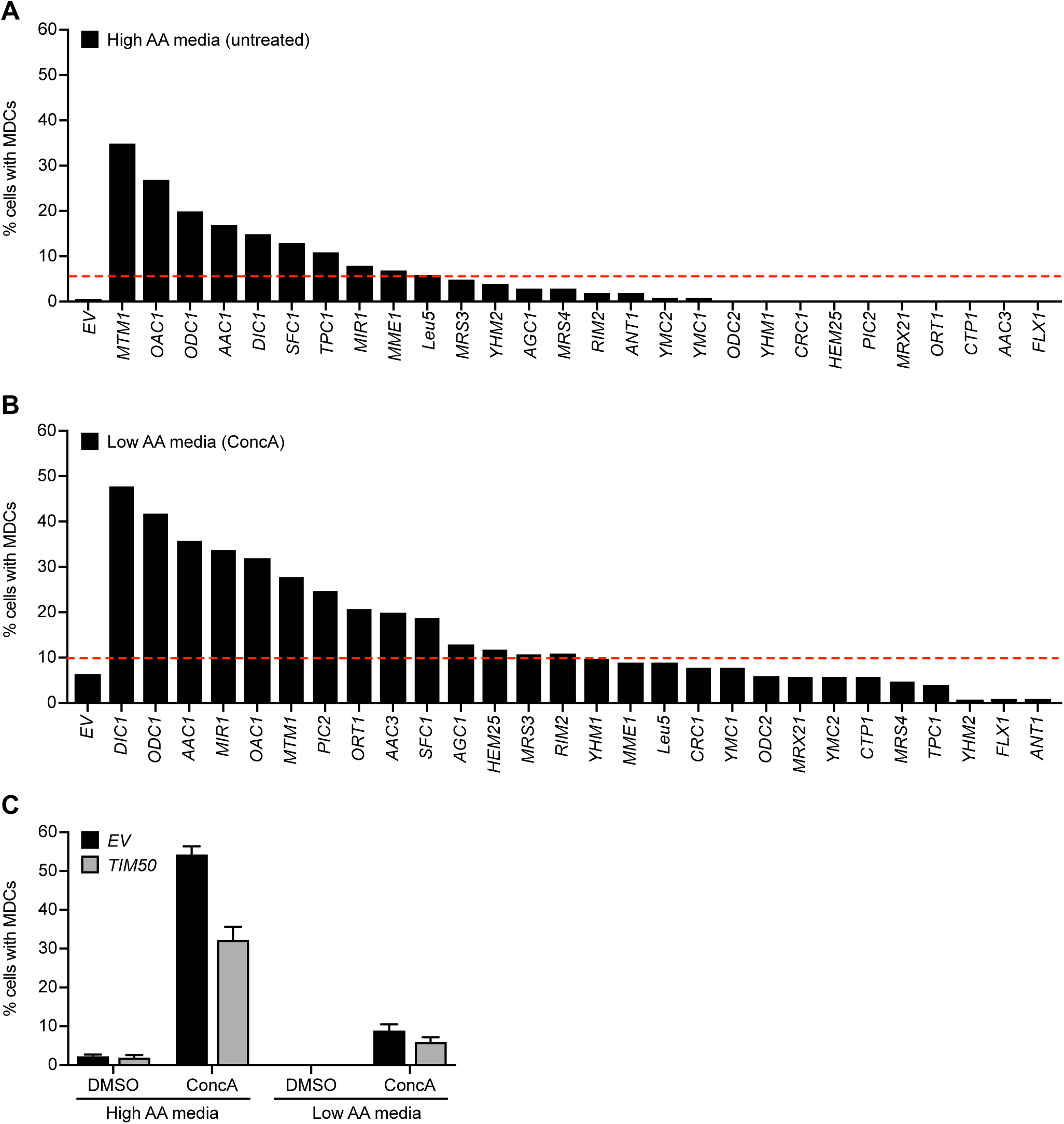
Elevated SLC25 Carrier Expression Activates MDC Formation, Related to Figure 6. (A) Quantification of MDC formation in cells expressing the indicated mitochondrial carrier in high amino acid media in absence of drug treatment. *n* = 100 cells. (B) Quantification of MDC formation in cells expressing the indicated mitochondrial carrier in low amino acid media in presence of ConcA. *n* = 100 cells. (C) Quantification of ConcA-induced MDC formation in *TIM50 over*expressing or *empty vector (EV)* control cells in high and low amino acid media. Error bars show mean ± SE of three replicates with *n* = 100 cells per replicate.

**Figure S7.**
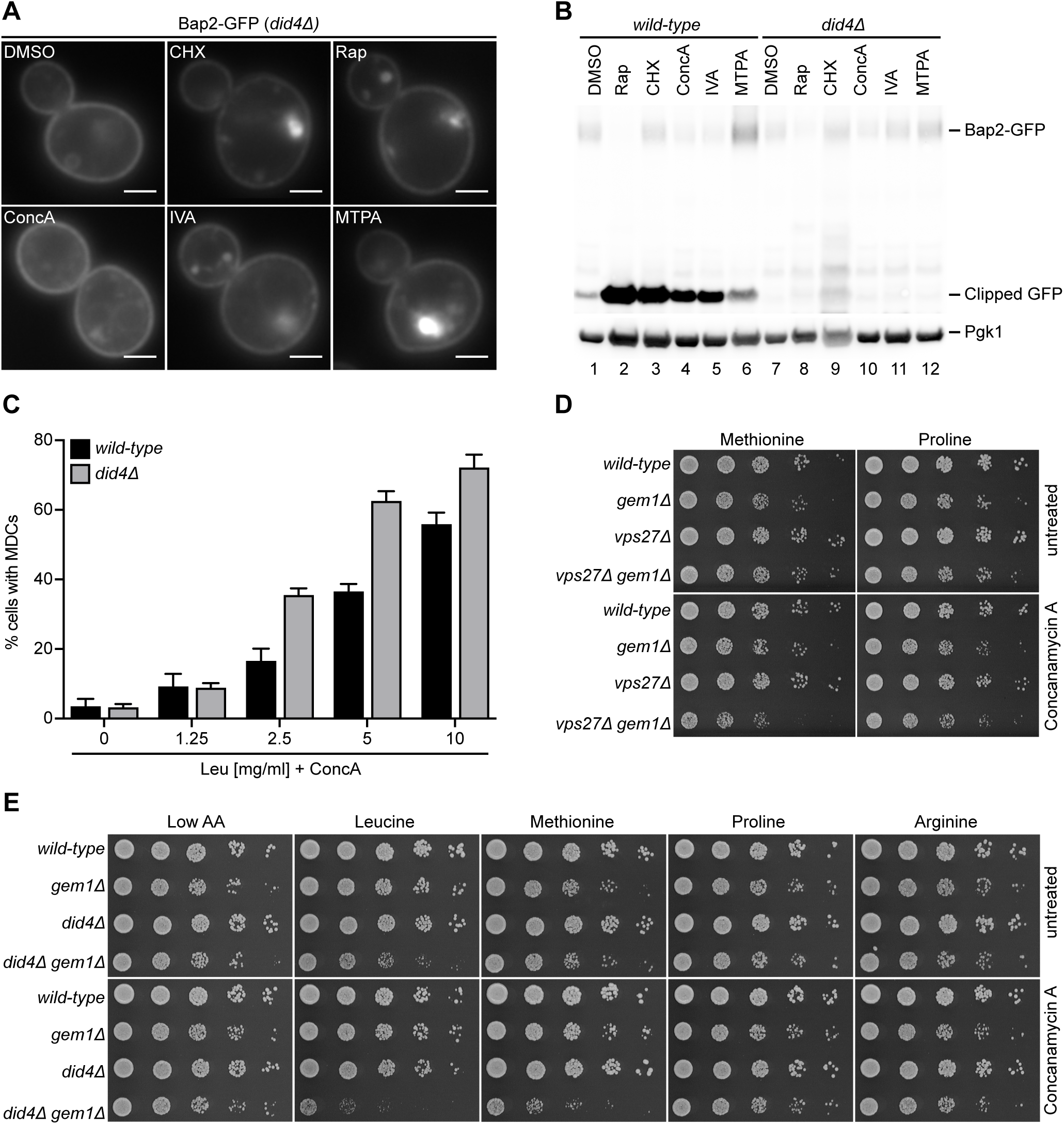
MDCs Cooperate with MVBs to Protect Cells from Amino Acid Toxicity, Related to Figure 7. (A) Widefield images showing stabilization of Bap2-GFP on the plasma membrane in rapamycin (Rap), cycloheximide (CHX), concanamycin A (ConcA), isovaleraldehyde (IVA) or 3-methylthiopropanal (MTPA) treated *did4Δ* cells. Bright structures resemble typical class E compartments found in ESCRT mutants. Scale bar = 2 µm. (B) Western blot analysis of Bap2-GFP clipping upon treatment with Rap, CHX, ConcA, IVA and MTPA in *wild-type* and *did4Δ* cells. Pgk1 – loading control. (C) Quantification of ConcA-induced MDC formation in low amino acid media supplemented with the indicated amount of leucine in *wild-type (WT)* and *did4Δ* cells. Error bars show mean ± SE of three replicates with *n* = 100 cells per replicate. (D) Growth of *wild-type*, *gem1Δ, vps27Δ* and *gem1Δvps27Δ* strains in presence and absence of ConcA on low amino acid media supplemented with 10mg/ml methionine or proline. (E) Growth of *wild-type*, *gem1Δ, did4Δ* and *gem1Δ did4Δ* strains in presence and absence of ConcA on low amino acid media supplemented with 10mg/ml leucine, methionine, proline or arginine.

## KEY RESOURCES TABLE

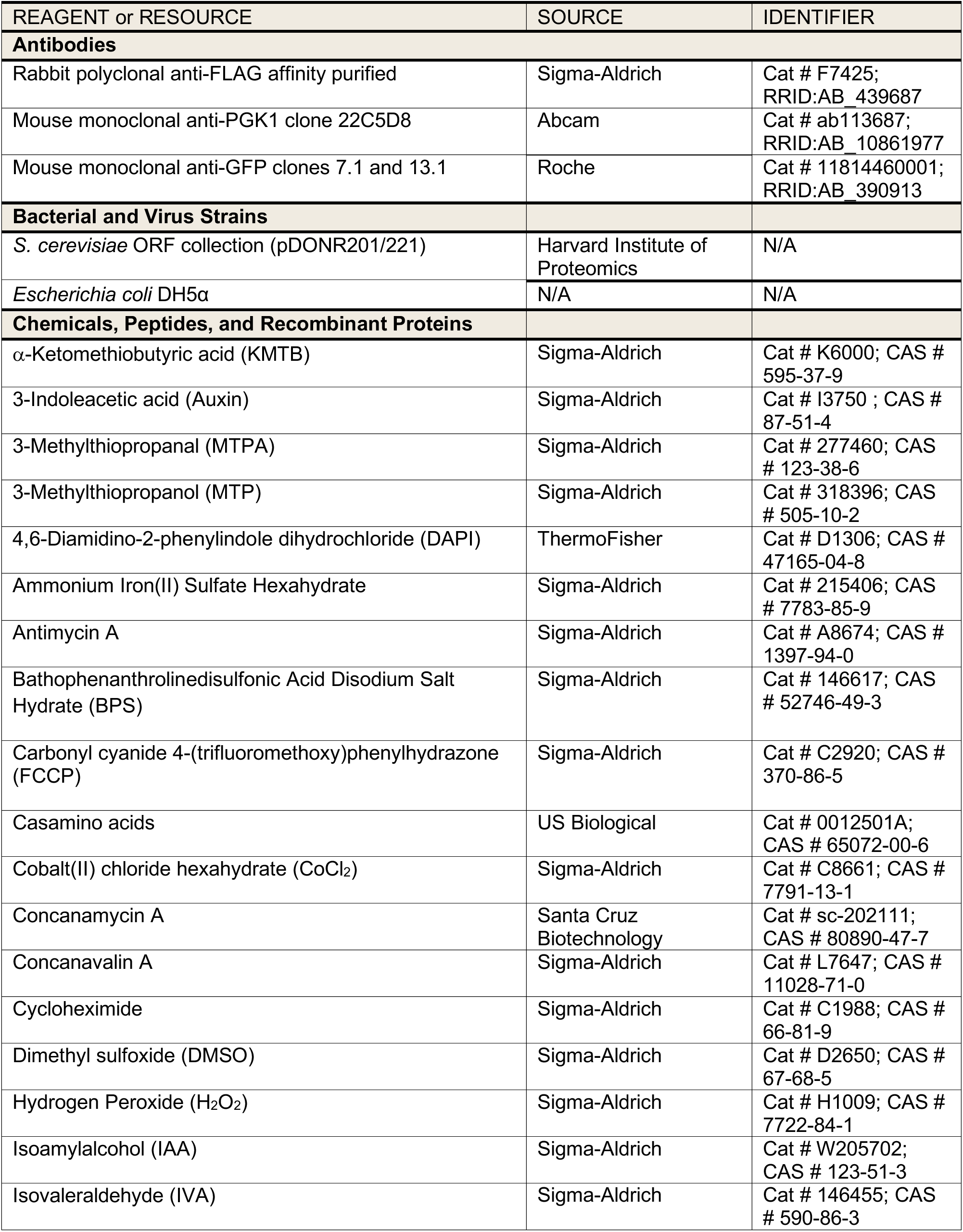
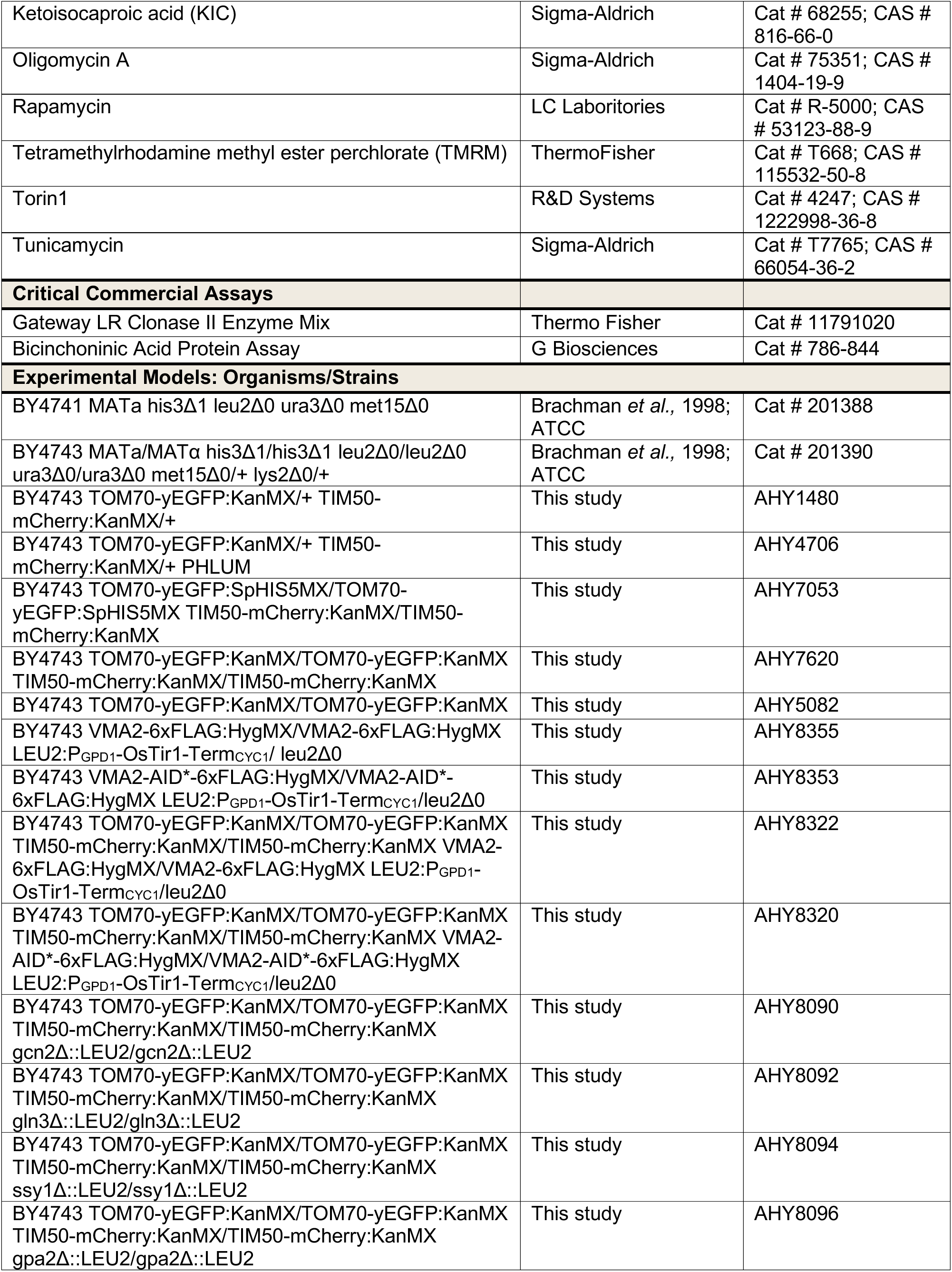
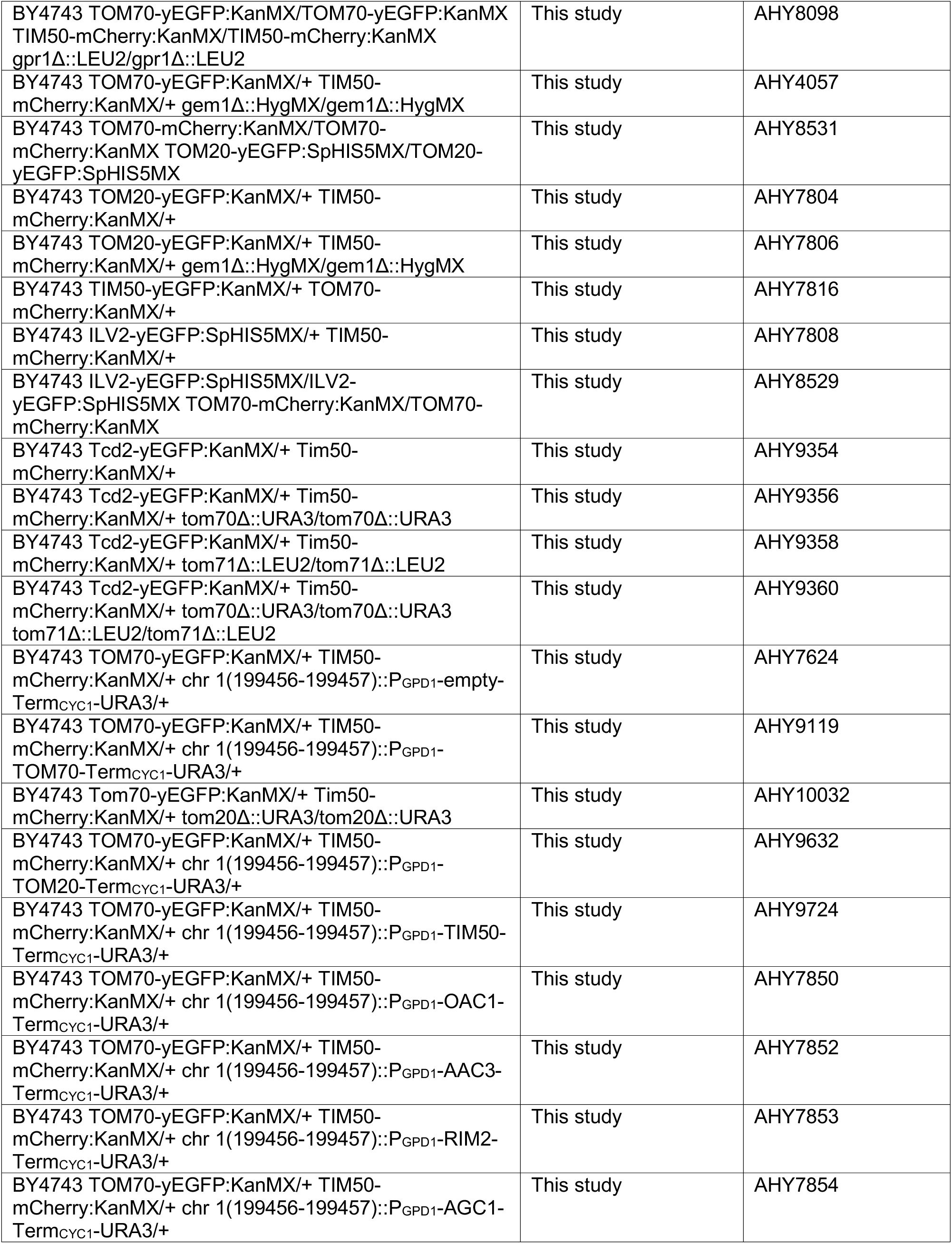
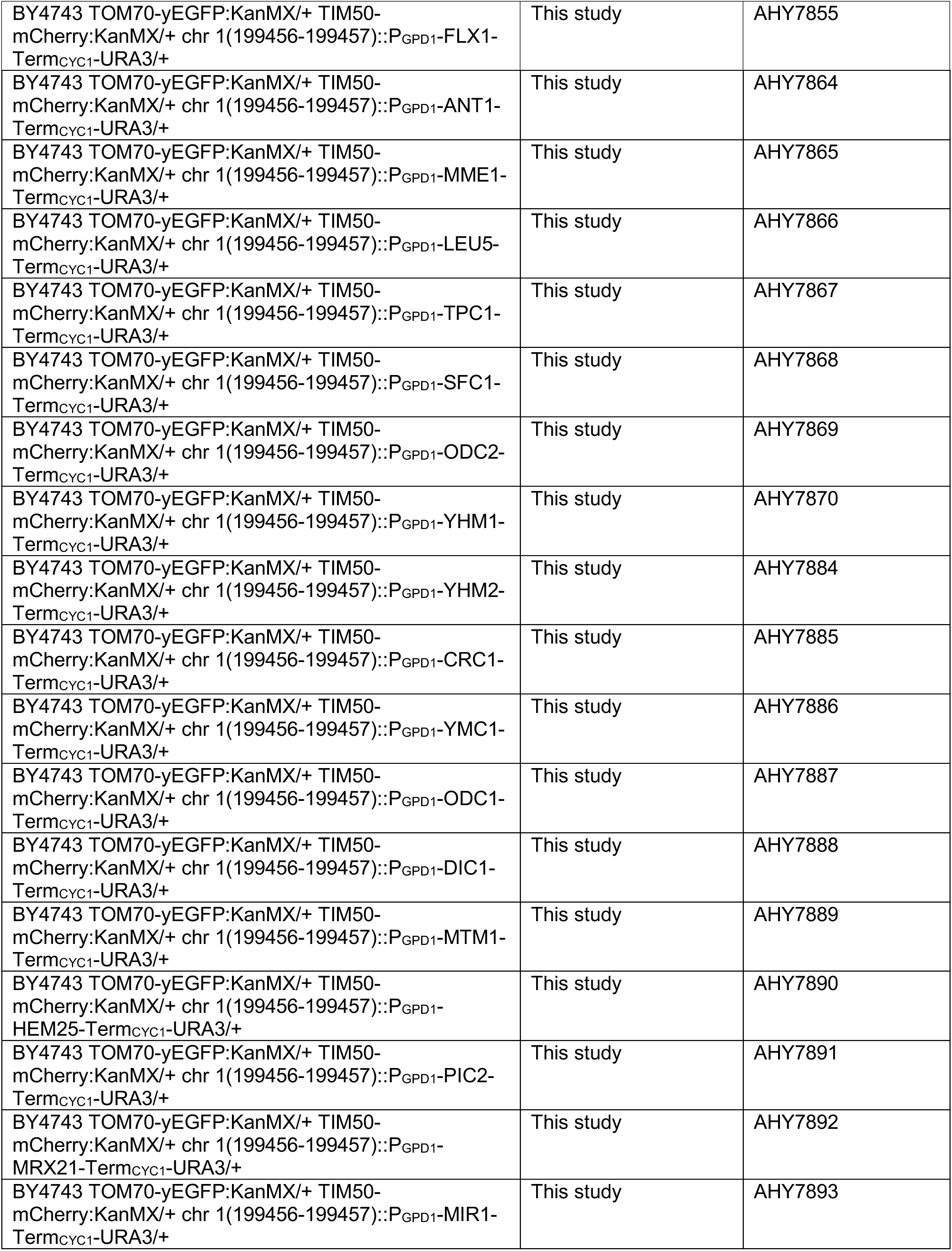
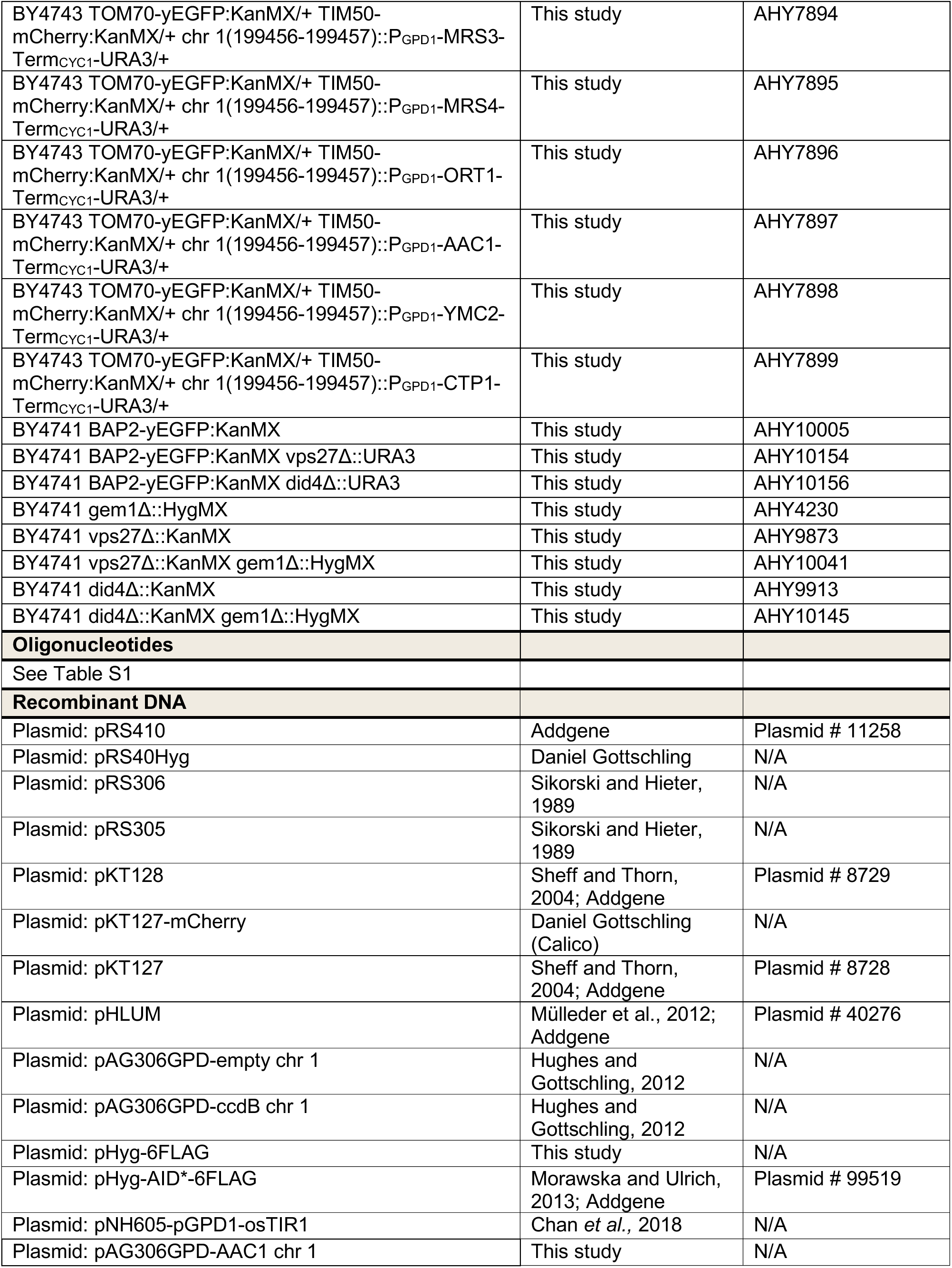
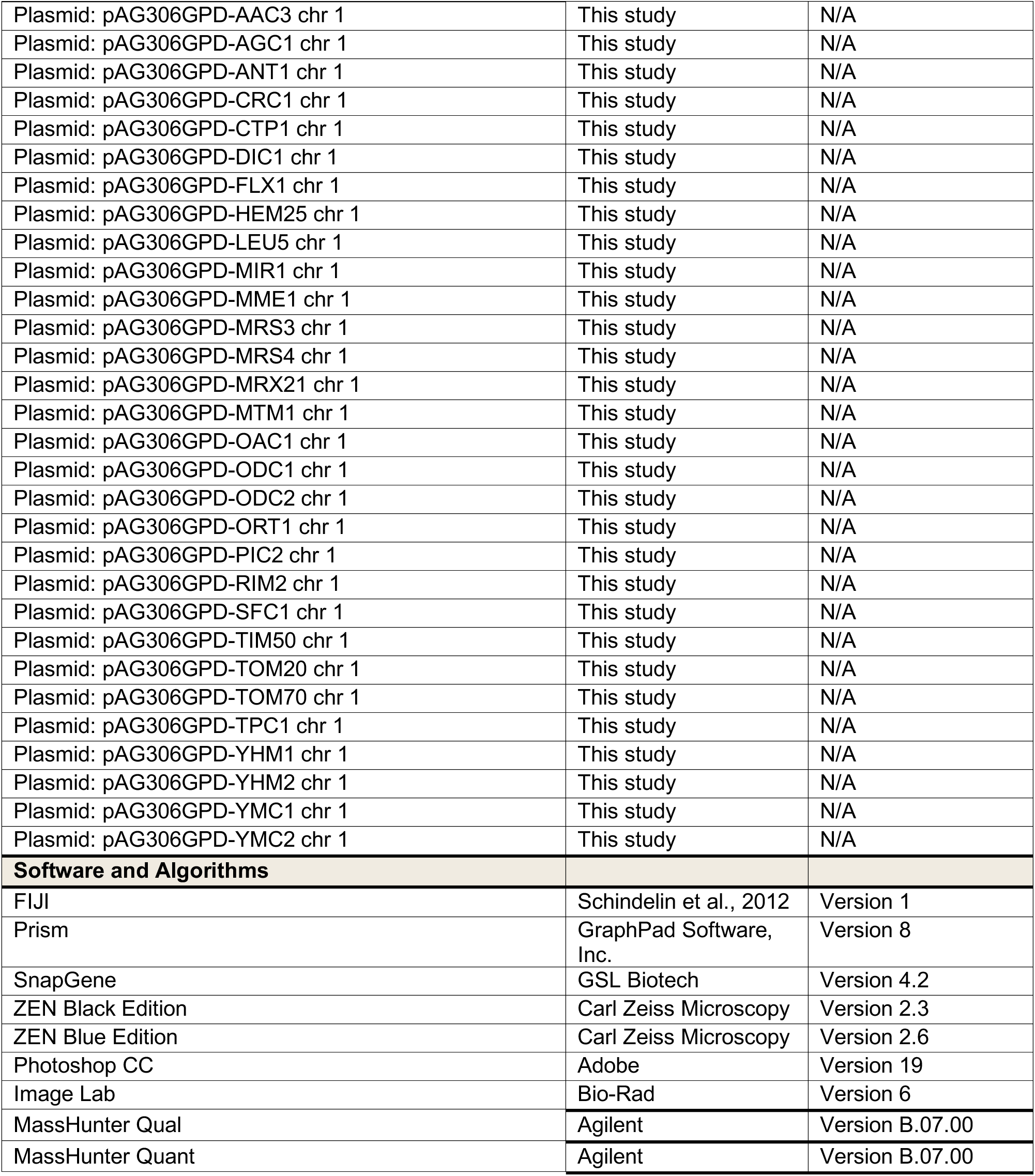

## SUPPLEMENTAL ITEM LEGENDS

**Video S1. Concanamycin A-Induced MDC Formation, Related to Figure 1**

An example of concanamycin A-induced MDC formation (from Figure 1G). Maximum intensity projection video of yeast expressing Tom70-GFP treated with concanamycin A. Images were taken every minute and are shown at 2 frames per second.

**Video S2. Dynamic MDC Behavior, Related to Figure 1**

An example of dynamic MDC behavior (from Figure S1E). Maximum intensity projection video of yeast expressing Tom70-GFP treated with concanamycin A. Images were taken every minute and are shown at 2 frames per second.

**Video S3. Cycloheximide-Induced MDC Formation, Related to Figure 2**

An example of cycloheximide-induced MDC formation (from Figure S2D). Maximum intensity projection video of yeast expressing Tom70-GFP treated with cycloheximide. Images were taken every minute and are shown at 2 frames per second.

**Video S4. Rapamycin-Induced MDC Formation, Related to Figure 4**

An example of rapamycin-induced MDC formation (from Figure S4C). Maximum intensity projection video of yeast expressing Tom70-GFP treated with rapamycin. Images were taken every minute and are shown at 2 frames per second.

**Table S1. Oligonucleotides Used in This Study, Related to Star Methods**

## CONTACT FOR REAGENT AND RESOURCE SHARING

Further information and requests for resources and reagents should be directed to and will be fulfilled by the Lead Contact, Adam Hughes. All unique/stable reagents generated in this study are available from the Lead Contact without restrictions.

## METHOD DETAILS

### Yeast Strains

All yeast strains are derivatives of *Saccharomyces cerevisiae* S288c (BY) (Brachmann et al., 1998). Strains expressing fluorescently tagged *TOM70*, *TIM50*, *TOM20*, *ILV2, TCD2, BAP2* and/or AID*-6xFLAG/6xFLAG tagged *VMA2* from their native loci were created by one step PCR-mediated C-terminal endogenous epitope tagging using standard techniques and oligo pairs listed in Table S1. Plasmid templates for fluorescent epitope tagging were from the pKT series of vectors (Sheff and Thorn, 2004). Plasmids used for AID*-6xFLAG/6xFLAG tagging and integration of GPD-OsTir1 into the *LEU2* locus are described below. Correct integrations were confirmed by a combination of colony PCR across the chromosomal insertion site and correctly localized expression of the fluorophore by microscopy. Deletion strains for *TOM70*, *TOM71*, *TOM20*, *GEM1, VPS27* and/or *DID4* were generated by one step PCR-mediated gene replacement with the indicated selection cassette using standard techniques and oligo pairs listed in Table S1. Plasmid templates for gene replacement were from the pRS series of vectors (Sikorski and Hieter, 1989). Correct insertion of the selection cassette into the target gene was confirmed by colony PCR across the chromosomal insertion site. Yeast strains constitutively expressing *TOM70, TOM20, TIM50* or the indicated mitochondrial carrier protein from the GPD promoter were generated by integration of the expression cassette into yeast chr I (199456-199457). Plasmids for integration of the GPD-driven expression cassette are described below. Correct insertion of the expression cassette into chr I was confirmed by colony PCR across the chromosomal insertion site.

*Wild-type* yeast strain AHY4706, which was rendered prototrophic with pHLUM (see below) to prevent complications caused by amino acid auxotrophies in the BY strain background, was used to quantify amino acid dependencies of MDC formation and for analysis of whole cell metabolite levels. *Wild-type* yeast strains AHY5082, AHY7053, AHY7620, AHY8529 and AHY8531 were used for super resolution and time-lapse imaging. *Wild-type* and *gem1Δ* yeast strains AHY4706, AHY4057, AHY7808, AHY7816, AHY7804, AHY7806 were used for quantification of MDC-dependent removal of proteins from mitochondria and protein enrichment in the MDC. A complete list of all strains used in this manuscript can be found in the Key Resources Table.

### Plasmids

Plasmids used in this study are listed in the Key Resources Table. pHLUM, a yeast plasmid expressing multiple auxotrophic marker genes from their endogenous promoters, was obtained from Addgene (#40276) (Mulleder et al., 2012). pHyg-AID*-6FLAG (Morawska and Ulrich, 2013) and pNH605-pGPD1-osTIR1 (Chan et al., 2018) were described previously. To integrate GPD1-osTIR1 into the *LEU2* locus, pNH605-pGPD1-osTIR1 was digested with Swa1. pHyg-6FLAG was generated by inserting 6FLAG amplified from pHyg-AID*-6FLAG into Kpn1/Xba1 digested pHyg-AID*-6FLAG. Plasmids for GPD-driven expression of *TOM70, TOM20, ILV2, TIM50* or the indicated mitochondrial carrier protein were generated by gateway mediated transfer of the corresponding ORF (Harvard Institute of Proteomics) from pDONR201/221 into pAG306GPD-ccdB chr 1 (Hughes and Gottschling, 2012) using Gateway™ LR Clonase™ II Enzyme mix (ThermoFisher) according to the manufacturer’s instructions. To integrate the resulting expression plasmid into yeast chr I (199456-199457), pAG306GPD-ORF chr 1 was digested with NotI. All insert sequences were verified by the University of Utah Sequencing Core.

### Yeast Cell Culture and Media

Yeast cells were grown exponentially for 15 hours at 30°C to a maximum density of 6×10^6^ cells/mL before the start of all experiments described in the paper, including MDC and spot assays. This period of overnight log-phase growth was carried out to ensure vacuolar and mitochondrial uniformity across the cell population and is essential for consistent MDC activation. Cells were cultured as indicated in the Main Text and Figure Legends in media containing high amino acids (1% yeast extract, 2% peptone, 0.005% adenine, 2% glucose) or low amino acids (0.67% yeast nitrogen base without amino acids, 2% glucose, supplemented nutrients 0.074 g/L each adenine, alanine, arginine, asparagine, aspartic acid, cysteine, glutamic acid, glutamine, glycine, histidine, myo-inositol, isoleucine, lysine, methionine, phenylalanine, proline, serine, threonine, tryptophan, tyrosine, uracil, valine, 0.369 g/L leucine, 0.007 g/L para-aminobenzoic acid). For growth in medium containing no amino acids (0.67% yeast nitrogen base without amino acids, 2% glucose), cells were cultured in low amino acid medium and then shifted to medium containing no amino acids at time of drug treatment. Where casamino acids (CasAA) were added to low or no amino acid media, CasAA were added at time of drug treatment to a final concentration of 2% w/v. For single amino acid re-addition experiments, individual amino acids were added to medium containing low amino acids at the time of drug treatment. All amino acids were added to a final concentration of 20 mg/mL, except indicated otherwise, with the exception of cysteine and tyrosine, which were added at final concentrations of 5 mg/mL and 1 mg/mL respectively, due to toxicity and/or solubility issues. Leucine and methionine catabolites were added at a final concentration of 10 μM at the time of drug treatment. Drugs were added to cultures at final concentrations of Concanamycin A (500nM), Cycloheximide (10 μg/mL), Rapamycin (250 nM), Torin1 (5 μM), Antimycin A (40 μM), FCCP (10 μM), Oligomycin (10 μM), H_2_O_2_ (10 μM), CoCl_2_ (1 mM), Tunicamycin (5 μg/ml) and BPS (250 μM). Iron was added to cultures as (NH_4_)_2_Fe(SO_4_) _2_(H_2_O)_6_ at a final concentration of 2 mM.

### Yeast MDC Assays

For yeast MDC assays, overnight log-phase cell cultures were grown in the presence of dimethyl sulfoxide (DMSO) or the indicated drug or metabolite for two hours. To acutely inhibit V-ATPase function by Auxin-inducible degradation of Vma2, overnight log-phase cell cultures were treated with 1 mM Auxin in high amino acid medium for the indicated times. For amino acid re-addition MDC assays, overnight log-phase cell cultures grown in low amino acid medium were shifted to low amino acid medium supplemented with the indicated amino acid and drug and grown for two hours. After incubation, cells were harvested by centrifugation, resuspended in imaging buffer (5% w/v Glucose, 10mM HEPES pH 7.6) and optical Z-sections of live yeast cells were acquired with an AxioImager M2 (Carl Zeiss) or, for super-resolution images, an Airyscan LSM800 (Carl Zeiss) or Airyscan LSM880 (Carl Zeiss). The percentage of cells with MDCs was quantified in each experiment at the two-hour time point from maximum intensity projected wide field images generated in ZEN (Carl Zeiss). MDCs were identified as Tom70-enriched, Tim50-negative structures of varying size and shape. In *tom70Δ* and *tom70Δ tom71Δ* strains, Tcd2, another MDC substrate (Hughes et al., 2016) was used to identify MDCs. A single focal plane is displayed for all yeast images with the exception of time-lapse images (see below).

### Yeast Time-Lapse Imaging

For time-lapse imaging of yeast treated with concanamycin A, overnight log-phase cultures grown in high amino acid medium were treated with concanamycin A for 30 minutes. Cells were harvested by centrifugation, resuspended in low amino acid medium plus Cas AA and concanamycin A, and pipetted into flow chamber slides as previously described (Fees et al., 2017). Briefly, flow chambers were made using standard microscope slides and coverslips. Strips of parafilm were used to seal a coverslip to a slide and created the walls of the chamber. Flow chambers were coated with concanavalin A prior to loading cells. Melted Vaseline was used to seal the chamber. MDC formation was imaged every minute for 120 minutes. For time-lapse imaging of yeast treated with cycloheximide, overnight log-phase cultures grown in high amino acid medium were treated with cycloheximide for two minutes. Cells were harvested by centrifugation, resuspended in low amino acid medium plus Cas AA and cycloheximide, and pipetted into flow chamber slides. MDC formation was imaged every minute for 120 minutes. For time-lapse imaging of yeast treated with rapamycin, overnight log-phase cultures grown in high amino acid medium were treated with rapamycin for 15 minutes. Cells were harvested by centrifugation, resuspended in low amino acid medium plus Cas AA and rapamycin, and pipetted into flow chamber slides. MDC formation was imaged every minute for 119 minutes. 300 nm optical Z-sections of live yeast cells were acquired with an Airyscan LSM880 in Airyscan fast mode. Time-lapse images and movies of yeast MDC formation show maximum intensity projections.

### Fluorescent Staining

In order to visualize mitochondrial DNA in yeast, cells were incubated with 2 μg/mL DAPI in high amino acid medium for 30 minutes at RT. To visualize mitochondrial membrane potential in yeast, cells were re-isolated by centrifugation, washed with imaging buffer and stained with 50 nM tetramethylrhodamine methyl ester (TMRM) for 15 minutes at RT. In case of concanamycin A treatment, media were supplemented with 2mM iron during the treatment period to prevent loss of the mitochondrial membrane potential as previously described (Hughes et al., 2020). To visualize vacuolar acidity in yeast, cells were re-isolated by centrifugation and stained with 200 μM Quinacrine in high amino acid medium supplemented with 50 mM HEPES pH 7.6 for 10 minutes at 30°C followed by a 5-minute incubation on ice. Prior to imaging, cells were washed twice with imaging buffer.

### Microscopy and image analysis

For quantification of MDC formation or fluorescence intensities, 200 nm optical Z-sections of live yeast cells were acquired with an AxioImager M2 (Carl Zeiss) equipped with an edge 4.2 CMOS camera (PCO) and 63× or 100× oil-immersion objectives (Carl Zeiss, Plan Apochromat, NA 1.4). Super resolution images showing MDC formation in live yeast cells were acquired with an LSM800 or LSM880 (Carl Zeiss) equipped with an Airyscan detector (Carl Zeiss) and 63× oil-immersion objective (Carl Zeiss, Plan Apochromat, NA 1.4). Widefield images were acquired with ZEN (Carl Zeiss), processed with Fiji (Schindelin et al., 2012), and represent single Z-sections. Super-resolution images were acquired with ZEN (Carl Zeiss), processed using the automated Airyscan processing algorithm in ZEN (Carl Zeiss) and Fiji, and represent single 200 nM Z-sections (with the exception of time-lapse images). Individual channels of all images were minimally adjusted in Fiji to match the fluorescence intensities between channels for better visualization. Line-scan analysis and measurements of MDC size were performed on non-adjusted, single Z-sections from super resolution images. To quantify the average MDC size, the diameter of spherical MDCs that had a visible lumen was measured using the line tool in Fiji. Tubular MDCs or smaller Tom70-enriched structures lacking Tim50 were excluded from the analysis since they likely represent broken down or growing MDCs. Fluorescence intensity analysis was performed on non-adjusted, maximum intensity projected wide-field images. To quantify the residual mitochondrial fluorescence upon MDC formation, the mean fluorescence intensity was measured along the entire mitochondrial tubule, excluding the MDC, using the freehand selection tool in Fiji and normalized to the untreated control sample. For analysis of fluorescence enrichment in the MDC, mean fluorescence intensities were measured in along the brightest part of the MDC using the freehand line tool in Fiji and normalized to the mean fluorescence intensity in the directly adjacent mitochondrial tubule.

### Yeast Growth Assays

To analyze growth of yeast cells on plates containing high levels of single amino acids, five-fold serial dilutions of over-night log-phase cultures grown in low amino acid medium were prepared in low amino acid medium and 3 µl of each dilution were spotted onto the agar medium (3% w/v agar) denoted in each Figure Legend. Total cells plated in each dilution spot were 5,000, 1,000, 200, 40, and 8.

### Protein Preparation and Immunoblotting

For western blot analysis of total protein levels, overnight log-phase cultures were treated as indicated and 5×10^7^ yeast cells were re-isolated by centrifugation, washed with ddH_2_O and incubated in 0.1 M NaOH for 5 min at RT. Subsequently, cells were re-isolated by centrifugation at 16,000 ×g for 10 minutes at 4°C and lysed for 5 minutes at 95°C in SDS-Lysis buffer (10mM Tris pH 6.8, 100 mM NaCl, 1 mM EDTA, 1mM EGTA, 1% (w/v) SDS). Upon lysis, protein concentrations were determined by bicinchoninic acid assay (G Biosciences) and samples were denatured in Laemmli buffer (63 mM Tris pH 6.8, 2% (w/v) SDS, 10% (v/v) glycerol, 1 mg/ml bromophenol blue, 1% (v/v) β-mercaptoethanol) for 5 minutes at 95°C. To separate proteins based on molecular weight, equal amounts of protein were subjected to SDS polyacrylamide gel electrophoresis and transferred to PVDF membrane (Millipore) by wet transfer. Nonspecific antibody binding was blocked by incubation with TBS containing 5% (w/v) dry milk (Sigma Aldrich) for one hour at RT. After incubation with the primary antibodies (listed in Key Resources Table) for two hours at RT or at 4°C overnight, membranes were washed five times with TBS and incubated with secondary antibody (goat-anti-rabbit/mouse HRP-conjugated,1:2000 in TBS + 5% dry milk, Sigma Aldrich) for 45 minutes at RT. Membranes were washed five times with TBS, enhanced chemiluminescence solution (Thermo Fisher) was applied and the antibody signal was detected with a BioRad Chemidoc MP system. All blots were exported as TIFFs using ImageLab 6.0 (BioRad) and cropped in Adobe Photoshop CC. Western blots show one representative blot from *n* = 3 replicates performed in parallel with the associated MDC assay.

### Extraction of Whole Cell Metabolites from Yeast

For analysis of whole cell metabolite levels, cells were grown exponentially in the indicated media for 15 hours to a maximum density of 6×10^6^ cells/mL and treated for two hours with the indicated drugs. 5 x 10^7^ total yeast cells were harvested by centrifugation, washed once with water, and cell pellets were shock frozen in liquid nitrogen. Whole cell metabolites were extracted from yeast cell pellets as previously described with slight modifications (Canelas et al., 2009). Briefly, the internal standard succinic-*d*_4_ acid (Sigma Aldrich 10907HD) at a final concentration of 1 μg/sample. Subsequently, 5 mL of boiling 75% EtOH were added to each cell pellet, followed by vortex mixing and incubation at 90°C for five minutes. Cell debris were removed by centrifugation and the supernatant was transferred to new tubes and dried *en vacuo*. Pooled quality control samples were made by removing a fraction of collected supernatant from each sample and process blanks were made using only extraction solvent and no cell culture pellet.

### GC-MS Analysis

Amino acid composition analysis was performed on an Agilent 7200 GC-QToF mass spectrometer fitted with an Agilent 7890 gas chromatograph and an Agilent 7693A automatic liquid sampler. Dried samples were suspended in 40 μL of 40 mg/mL O-methoxylamine hydrochloride (MP Biomedicals) in dry pyridine and incubated for one hour at 30°C. 25 μL of this solution was added to auto sampler vials. 60 μL of N-methyl-N-trimethylsilyltrifluoracetamide (Pierce) was added automatically via the auto sampler and incubated for 30 minutes at 37°C with shaking. After incubation, 1 μL of the prepared sample was injected into the gas chromatograph inlet in the split mode with the inlet temperature held at 250°C. A 25:1 split ratio was used for analysis. The gas chromatograph had an initial temperature of 60°C for one minute followed by a 10°C/min ramp to 325°C and a hold time of 2 minutes. A 30-meter Agilent Zorbax DB-5MS with 10m Duraguard capillary column was employed for chromatographic separation. Helium was used as the carrier gas at a rate of 1 mL/min.

Data was collected using MassHunter software (Agilent). Metabolites were identified and their peak area was recorded using MassHunter Quant. Metabolite identity was established using a combination of an in-house metabolite library developed using pure purchased standards, the NIST library and the Fiehn Library. Values for each metabolite were normalized to the internal standard in each sample and are displayed as fold change compared to the control sample. All error bars show the mean ± standard error from three biological replicates analyzed in the same GC-MS run.

### Quantification and Statistical Analysis

All experiments were repeated at least three times. All attempts at replication were successful. Sample sizes were as large as possible to be representative, but of a manageable size for quantifications. Specifically, for yeast MDC assays, *N* = three replicates, with *n* = 100 cells for each replicate, for quantification of MDC size, *N* = 30 - 50 MDCs from *N* = 3 - 5 experiments and for fluorescence intensity analysis *N* = 45 cells from *N* = three replicates with *N* = 15 cells per replicate. No data were excluded from the analyses. No randomization or blinding was used as all experiments were performed with defined laboratory reagents and yeast strains of known genotypes.

